# Molecular mechanisms of anti-diabetes effects of *Mormodica charantia* L. Ethanolic leaf extract on type 2 diabetes in strains of *Drosophila melanogaster* meigen, 1838

**DOI:** 10.1101/2025.08.11.669659

**Authors:** M A Agi, R Abdulazeez, T N Dakup, A A Nabilah, A H Omodele, Y Wada, D M Shehu

## Abstract

Type 2 diabetes (T2D) is a chronic metabolic disorder characterized by reduced insulin sensitivity and poor glucose utilization. This study examined the anti-diabetic potential and molecular mechanisms of *Mormodica charantia* (MC) ethanolic leaf extract in strains of *Drosophila melanogaster*. T2D was induced using a high sucrose diet (2.5 g/10 g), followed by co-administration of *M. charantia* (MC) extract (100 mg, 150 mg, and 200 mg) and 16 mg of metformin for ten days. MC at 200 mg significantly improved eclosion rates and starvation resistance in the Harwich strain, while 150 mg showed the strongest hypoglycemic effect in the indigenous NgD3 strain. The extract alleviated heat stress and reversed sucrose-induced Malpighian tubule dysfunction, (p<0.001), which elevated creatinine (1.92 mg/dl and 1.59 mg/dl) both sexes and sodium levels (1.67 mg/dl), particularly in male flies. At 150 mg, effectively reduced creatinine and urea levels (1.68 µ/l). The 100 mg increased ALP levels (9.09 µ/l). At 200mg, *M. charantia,* significantly upregulated *CG4607* expression (0.5656) in Harwich strain, while causing a significant downregulation (-5.79586) in NgD3 strain. Squalene demonstrated the highest binding affinity (-7.5 kcal/mol). Single Nucleotide Polymorphism analysis revealed strain-specific genetic responses, and phylogenetic analysis showed three distinct clusters. These findings suggest MC extract modulates physiological and molecular responses in insulin-resistant Drosophila, highlighting its potential as a plant-based anti-diabetic therapy.

## INTRODUCTION

Diabetes is a chronic metabolic disorder characterized by elevated blood glucose levels (CDC, 2018). It arises either due to insufficient insulin production by the pancreas or the body’s inability to effectively utilize the insulin produced (WHO, 2023). Managing diabetes requires continuous medical care and multifactorial risk-reduction strategies beyond mere glucose control (ADA, 2023). Type 2 diabetes mellitus (T2DM) is one of the leading public health problems worldwide, causing high morbidity and mortality rates among different age groups (Alame *et al*., 2020: Ibrahim *et al*., 2023). An alarming increase in the global prevalence of diabetes has been reported from 422 million in 2014 to 463 million in 2023, with a projected increase of 783 million in 2045, with approximately 80% of this increase occurring in developing countries (IDF, 2021: Zanzabil *et al*., 2023). Reduced insulin sensitivity, poor glucose utilization, hyperinsulinemia, and aberrant lipid build up are common characteristics of insulin resistance (IR), which is a critical pathogenic factor in several metabolic disorders, including type 2 diabetes (T2D) (Lee *et al*., 2022; Zhao *et al*., 2023). Like obesity and type 2 diabetes, IR affects nearly half of the world’s population (Chooi *et al*., 2019; Sun *et al*., 2022) and 3.6 million people in Nigeria (International Diabetes Federation, 2021). The modern diabetogenic lifestyle, which includes the consumption of extra energy, primarily carbohydrates, is one of the risk factors most closely linked to the present T2D epidemic (Hosseini *et al*., 2022).

One plant that has gained substantial attention in this regard is *M. charantia* is commonly referred to as “bitter melon or bitter gourd” in English, “Garhuni” in Hausa, “Ejirin” in Yoruba and “Kakayi or Singha” in Igbo (Aminah and Anna, 2011). The plant belongs a member of the Cucurbitaceae family and is widely distributed across tropical and subtropical regions such as India, Malaya, China, Thailand, Japan, Singapore, Vietnam, Amazon, East Africa, Brazil, China, Colombia, Cuba, Ghana, Nigeria Haiti, India, Mexico, Malaya, New Zealand, Nicaragua, Panama, Middle East, Central and South America (Shan *et al*., 2012; Shuo *et al*., 2017). Bioactive compounds like charantin, vicine, and polypeptide-p present in bitter melon can enhance insulin secretion and improve glucose tolerance, making it a popular natural remedy for diabetes management (Richter *et al*., 2023). The seeds and leaves of the plant are also used in traditional remedies to support overall health and manage blood sugar levels (Hussain *et al*., 2024).

**Plate I:**
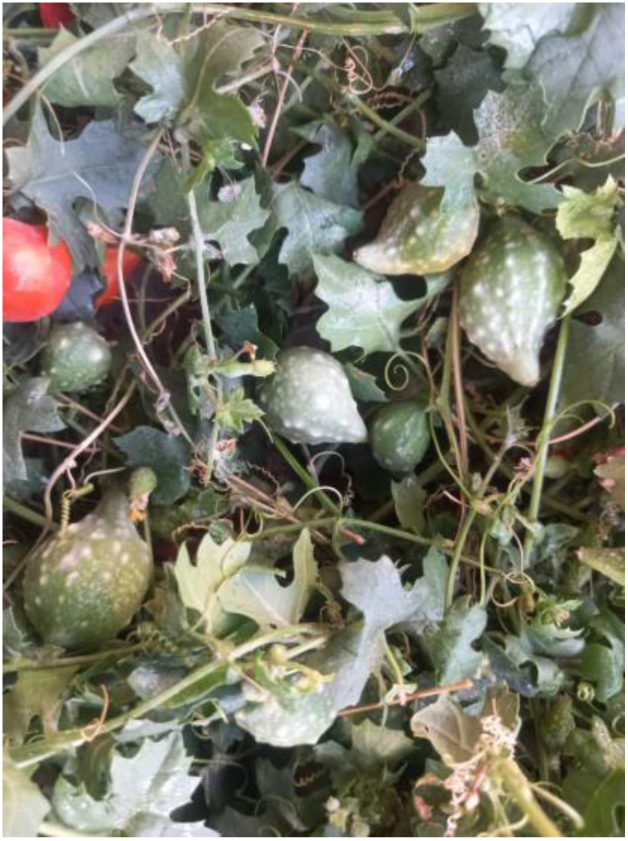
Mormodica charantia.

Although the identification of promising plant-based therapies is crucial, their efficacy and underlying mechanisms must be validated using suitable model organisms. One such model is Drosophila melanogaster, commonly known as the vinegar fruit-fly, which has gained prominence in biomedical research due to its high genetic similarity to humans (Omale et al., 2021). Although the insect body plan is simpler than that of mammals, the fruit flies shares anatomical and physiological analogs to mammalian organs such as the heart, lung, kidney, liver, gut and gonads (Staats et al., 2018). The fruit fly possesses a central and peripheral nervous system, produces gastrointestinal and sex hormones, insulin-like peptides, has conserved insulin signalling pathways, and exhibits about 75% homology with human disease-related genes (Adams et al., 2000; Staats et al., 2018). Key discoveries in classical genetics, including inheritance and gene function, have been made using Drosophila (Ashburner and Bergman, 2017; Yamaguchi and Yoshida, 2018). Advances in molecular biology have enabled the study of gene expression, development, and behaviour in this model organism.

## MATERIALS AND METHODS

### Study Area and Research Location

The study was conducted at the Drosophila and Neurogenetics Laboratory, Venom and Natural Toxins Research Centre (VANTRC), Ahmadu Bello University, Zaria (Latitude: 11° 8ʹ 18.938ʹʹ N and Longitude: 7° 38ʹ 43.052ʹʹ E), a suburb of Zaria in Kaduna State, Nigeria. Samaru, where the university is located, lies at Latitude: 11° 9ʹ14.07″ N. and Longitude: 7° 37ʹ 22.25″ E.

### Culture of *D. melanogaster* Strains

Two strains of *Drosophila melanogaster*, Harwich and Indigeneous (Ngd3), were obtained from Drosophila and Neurogenetics Laboratory, Department of Zoology, A.B.U. Zaria and maintained at the Drosophila and Neurogenetics Laboratory, VANTRC.

### Meal Preparation

Cornmeal was prepared according to the method described by Markow and O’Gardy (2006). A total of 850 ml of distilled water was measured 700 ml was transferred into a pot and brought to a boil, while 150 ml was used to dissolve 50g of corn flour. Separately, 10g of yeast was dissolved in a portion of the boiled water, and 8g of Agar-Agar was added into the boiling water, stirring periodically to avoid lumps. After cooking for 10 minutes, the dissolved corn flour solution was added, cooked for another 10 minutes, followed by the yeast solution left to cook for another 15 minutes. The mixture was allowed to cool slightly before adding 1g of Nipagin dissolved in 5 ml ethanol.

### Collection of *M. charantia*

Fresh vegetative parts of *M. charantia* were collected from the wild in Gashua, Bade Local Government area, Yobe State (Latitude: 12° 51’ 41.33″ N and Longitude: 11° 2’ 45.82″ E) early in the morning (7:00 am) (Figure 3.1). The samples were placed in appropriately labelled polythene bags, and transported to the Herbarium of the Department of Botany, A.B.U. Zaria for authentication using reference keys. The plant was assigned a voucher number: *ABU01597*.

### Induction of diabetes in flies

To induce type 2 diabetes in the flies, sucrose (2.5 g sucrose/10 g diet) was added to the regular fly diet, and all the standard fly food ingredients were kept constant. Flies were observed after ten days for symptoms of diabetes, which included delayed egg production, decreased body size for both larvae (L3) and adult flies and decreased locomotor activities, as described by Omale *et al*. (2021).

### Starvation Assay

Newly eclosed flies were anaesthetize using Ice-block of various treatment group. Flies are sorted into sexes and 10 flies were placed in each vial containing 1% Agar agar. Vial were kept at room temperature. Number of dead and alive were recorded at 12 hours intervals until all the flies are dead. (Rion and Kawecki, 2007).

### Eclosion Assay

An adult flies were placed on an egg laying cage for the eggs plates to be incubated at 25°C for 16-18 hours with a lid placed on it. The embryos are aged for 24-30 hours to establish plate full of 1^st^ instar (L1) larva. A 20 L1 was transferred into the medium of interest using paint brush after suspending medium in solution of sucrose. Development of L1 was recorded on the 11^th^ day of experimental set up, all flies that are eclosed are counted and recorded, dead or alive. Flies are anaesthetized with ice block and count by direct inspection (Rand *et al*., 2014).

### Heat Stress Tolerance Assay

Flies were transferred into empty vials (5 of each sex) allowed to acclimate, and placed in water bath at 35 °C for 60 minutes. The were transferred flies into their respective food vials immediately after administering the heat shock, for recovery. Mortality or immobilization was assessed immediately and at 24 hours post heat shock.

## BIOCHEMICAL ASSAY

### Creatinine assay

The level of Creatinine was determined using Jaffe’s method by mixing 0.5 µL of the fly’s homogenate with picric acid and sodium hydroxide solution. The sample was measure with spectrophotometer at 520 nm and absorbance was compare to the standard solution (Larsen, 1972).

### Urea assay

The level of Uric acid was determined by mixing 0.5 µL of the fly’s homogenate with diacetyl monoxime, sulfuric acid, Thiosemicarbazid and a catalase (Ferric ions). The sample was measure with spectrophotometer at 520 nm and absorbance was compare to the standard uric acid solution (Fawcett and Scott, 1960).

### Sodium assay

The level of sodium was determined using Flame Photometry Method by mixing 0.5 µL of the fly’s homogenate and diluted with 0.1% Lanthanum solution The sample was measure with spectrophotometer at 589 nm and absorbance was compare to the standard sodium solution (Bowers and Transue, 1956).

### Calcium assay

The level of calcium was determined using O-Cresolphthalein Complexone Method by mixing 0.5 µL of the fly’s homogenate and mix with o-Cresolphthalein complexone reagent, alkaline buffer and 8-hydroxyquinoline was added to the mixture and allow to react at a at room temperature (35 °C). The sample was measure spectrophotometrically at 570 nm. (Gitelman, 1967)

### Alkaline phosphatase (ALP) assay

The level of ALP was determined by mixing 1 µL of the fly’s homogenate and incubate at 37 °C for 20 minutes and mixed with p-nitrophenol and phosphate. The stop solution was add to halt the enzyme activity and stabilize the colour, the sample absorbance was measured at 405 nm using a spectrophotometer (Bessey *et al*., 1946).

### Alanine aminotransferase (ALT) assay

The level of ALT was determined by mixing 1 µL of the fly’s homogenate mixed with L-alanine and 2-oxoglutarate incubate at 37 °C for 30 minutes and mixed with DNPH to the reaction mixture to react with the pyruvate formed. The mixture was allowed for 20 minutes at room temperature. And NaOH was added to alkalize the solution and develop the colour. The sample absorbance was measured at 505 nm using a spectrophotometer (Reitman and Frankel, 1957).

### Aspartate aminotransferase (AST) assay

The level of ALT was determined by mixing 1 µL of the fly’s homogenate mixed with L-alanine and 2-oxoglutarate incubate at 37 °C for 30 minutes and mixed with DNPH to the reaction mixture to react with the pyruvate formed. The mixture was allowed for 20 minutes at room temperature. And NaOH was added to alkalize the solution and develop the colour. The sample absorbance was measured at 505 nm using a spectrophotometer (Reitman and Frankel. 1957).

### Malondialdehyde (MDA) Assay

The level of MDA was determined by mixing 1 µL of the fly’s homogenate mixed the with TCA, TBA, and HCl in a test tube. And was incubation at 95 °C for 60 minutes in a water bath, the sample was allowed to cool at room temperature. The sample absorbance was measured at 532 nm using a spectrophotometer. The MDA concentration was determined using the molar extinction coefficient of the MDA-TBA complex (Ohkawa *et al*., 1979).

### Catalase (CAT) assay

Catalase activity was measured by the hydrogen peroxide decomposition method as described by Aebi (1984). The fly homogenates was prepared in phosphate buffer (pH 7.0). The reaction was initiated by adding hydrogen peroxide (H₂O₂) to the sample, and the decomposition of H₂O₂ was monitored by measuring the decrease in absorbance at 240 nm for 1 - 3 minutes using a spectrophotometer. Catalase activity was determined based on the rate of decrease in absorbance.

### Superoxide Dismutase (SOD) Assay

Superoxide dismutase activity was assayed using the nitroblue tetrazolium (NBT) reduction method, following the protocol of Beauchamp and Fridovich (1971). Tissue homogenates were prepared in phosphate buffer (pH 7.8). The reaction mixture included NBT and xanthine, with xanthine oxidase added to generate superoxide radicals. After incubating the mixture at 25 °C for 10 minutes, the absorbance was measured at 560 nm. SOD activity was determined by the enzyme’s ability to inhibit the reduction of NBT to blue formazan.

### Glutathione (GSH) Assay

Glutathione levels were quantified using Ellman’s reagent (5,5’-dithiobis (2-nitrobenzoic acid), DTNB) according to the method of Ellman (1959), fly homogenates was prepared in phosphate buffer (pH 7.4), and Ellman’s reagent was added to the samples. The mixture was incubated at room temperature for 10 minutes. Absorbance of the yellow-colored product was measured at 412 nm, and GSH concentration was calculated using the molar extinction coefficient of the TNB complex.

### Evaluation of Gene Expression Patterns

Total RNA was extracted from homogenates of 10–15 flies using *Accurep*® Universal RNA Extraction Kit (K-3140, K-3141). The flies were homogenized, and 500 µL of RB buffer plus 5 µL of mercaptoethanol were added. The mixture was centrifuged at 13,000 rpm for 3min, and the supernatant was mixed with 200 µl of ethanol and passed through a binding column. To the column, 400 µL of RB Buffer was added to the cell pellet and mixed by vortex followed by the addition of 300 µL of ethanol (80%) and immediate mixing. The sample was then transferred to a binding column and centrifuged at 14,000 rpm for 20 sec. This was followed by a series of washing steps using 700 µL of RWA 1 and 500 µL of RWA2 buffers, with centrifugation 14,000 rpm for 20 seconds. After washing, the column was centrifuged at 14,000 rpm for 1 min to remove residual ethanol. Finally, the RNA was eluted by adding 50-200 µLof ER Buffer to the binding column, incubated for 1 min at room temperature, and centrifuged at 10,000 rpm for 1 min.

### RNA Reverse Transcriptase Reaction to cDNA

The *Tubulin* gene, which is a housekeeping gene was used was required to normalize the relative expression level of each gene. RNA was reverse-transcribed into cDNA using Ready Mix Taq and a three-step thermal cycle 94 °C for -5 min; 94 °C for 30 Sec, 54 °C for 30 Sec, 72 °C for 30 Sec, repeated for 35 cycles and 72 °C - for 5 min.

#### 3.9.6.2 Quantitative Real Time Polymerase Chain Reaction

Quantitative Real-Time Polymerase Chain Reaction (qPCR) was conducted using a TIANLONG^®^ Real-Time PCR system. RNA from 10–15 homogenized flies was used. Amplification was performed to quantify gene expression levels.

#### 3.9.6.3 The knockdown efficiency

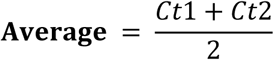

### In Silico Study

Molecular docking was conducted as described by Aminu *et al*., (2022). The full coding sequence of the *Drosophila* glucose transporter gene CG4607 was retrieved from NCBI. The structures of the *CG4607* protein was modeled using homology modeling tool, because the 3D structure is currently not available on the Protein DataBase (PDB). Homology modelling was performed using the SWISS-MODEL server with an AlphaFold template (https://swissmodel.expasy.org/). The resulting PDB file was validated via SAVES v6.0, including a Ramachandran plot. The 3D structures of bioactive compounds identified identified by GC-MS were obtained from the PubChem database (https://pubchem.ncbi.nlm.nih.gov/) and converted from SDF to PDB format using Open Babel GUI. Both ligands protein were prepared in Autodock 4.2, converted to PDBQT format, and docking was conducted using AutoDock Vina The grid size of (x = 68.00, y = 44.00, and z = 68.00) and center (x = -10.056, y = 3.45 and z = 3.339) covered the entire protein. The binding affinities were recorded, and the top two ligands were visualized Biovia Discovery Studio.

#### Evaluation of Genetic Variation

Genomic DNA was extracted from pools of 10-15 flies using AccuPower® 2X GreenStarTM qPCR MasterMix Kit (BIONEER Corporation) following the manufacturers’s protocol.

Flies were washed with distilled water homogenized using a squashing rod in lysis tubes. A total of 750 µL lysis solution was added, incubated for 1 hour, vortexed for 30 seconds, followed by the addition of 50 µL ethanol. After 10 sec seconds of vortexing, the mixture was centrifuged at 12,000 rpm for 30 seconds. The debris (supernatant) was transferred into the binding column, centrifuged for 1 minute, and the flow-through was discarded.

Next, 400 µL of inhibitors remover buffer was added and centrifuged for 1 minute. The flow-through was discarded. Then, 400 µL and 300 µL of deionized buffer (Buffer 2) were sequentially added and centrifuged at 12,000 rpm for 1 minute each. An additional centrifugeation for 1 minute at 12,000 rpm, ensured ethanol removal. DNA was eluted with 50 µL elution buffer after incubation at room temperature for 5 minutes and cold centrifuged (TOMY NTX-500) at 12,000 rpm for 2 minutes. DNA was stored for downstream applications.

#### Gene Amplification by PCR and Agarose Gel Electrophoresis

Polymerase chain reaction (PCR) was carried out to amplify *Drosophila* Glucose transporter (*CG4607*). A mix of 2 µL of primer and 6 µL distilled water was prepared from which 8 µL was combined with 2 µL DNA and added to a PCR tube containing the premix. The amplicons were resolved on 1.5 % agarose gel prepared in 1X TAE buffer, microwaved for 2 ½ minutes. Bands were visualized and documented using a phone camera.

#### DNA Sequencing

Sequencing was conducted at Inqaba Biotechnical Industries, South Africa. Sequences were analyzed for single nucleotide polymorphisms (SNPs) between strains to evaluate the genetic variations (Francis *et al*., 2021).

#### Data Analyses

Glucose and triglyceride levels were compared across groups using two-way ANOVA, and *Tukey’s post hoc* test. Geotaxis assay across groups was also analyzed using two-way ANOVA, and *Tukey’s post hoc* test Kaplan–Meier’s method analysis and the log-rank test were used for heat tolerance curve survival rates. and log rank test to check the homogeneity. The knockdown efficiency was calculated using delta-delta Ct method (Microsoft Excel and GraphPad Prism V10) and the Ct values were graphed. Significance between the control and knockdown transcript was calculated using Student’s t-test (p<0.001). Genetic diversity was evaluated through BLAST, multiple sequence alignment, construction using MEGAX11.

## RESULTS

### Male Starvation resistance of Harwich strain *D. melanogaster* fed with metformin and *M. charantia*

The mean starvation resistance of Harwich strain *D. melanogaster* co-exposed to high sucrose, M. charantia and Metformin is shows in (Figure 1a). Which indicate variation in starvation resistance for the control and treated group. For males, Negative and positive control group had the mean hours of flies survived for (70.7 and 92 hours) respectively, Metformin had the mean hours of flies that survived (74.7). Among the treatment group 100 mg had mean hours, survived for (69.3), 150 mg had (89.3) and 200 mg had mean hours survived for (72).

**Figure 1a:**
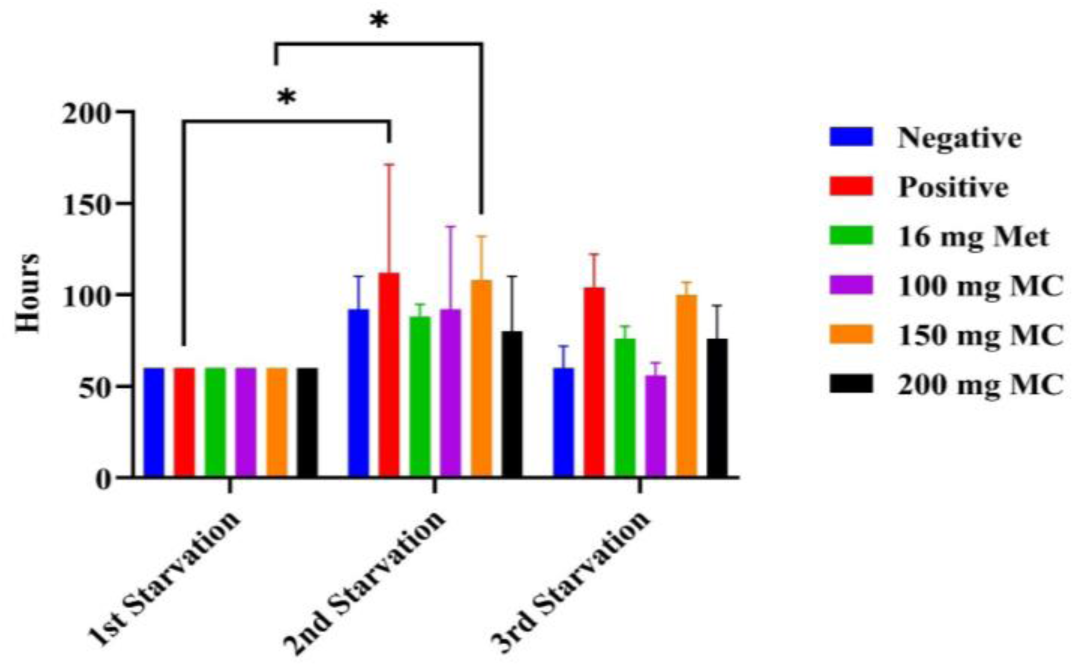
Males starvation resistance of Harwich strain *D. melanogaster* induced with Type II diabetes fed with metformin and MC.

**Figure 1b:**
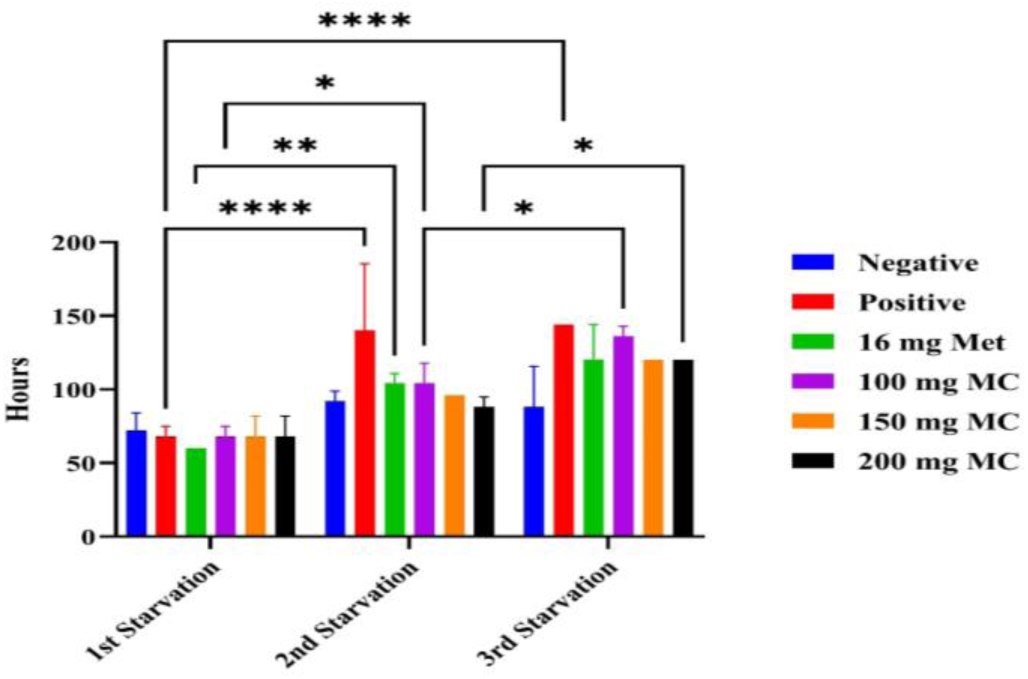
Females starvation resistance of Harwich strain *D. melanogaster* induced with Type II diabetes fed with metformin and MC.

**Figure 1c:**
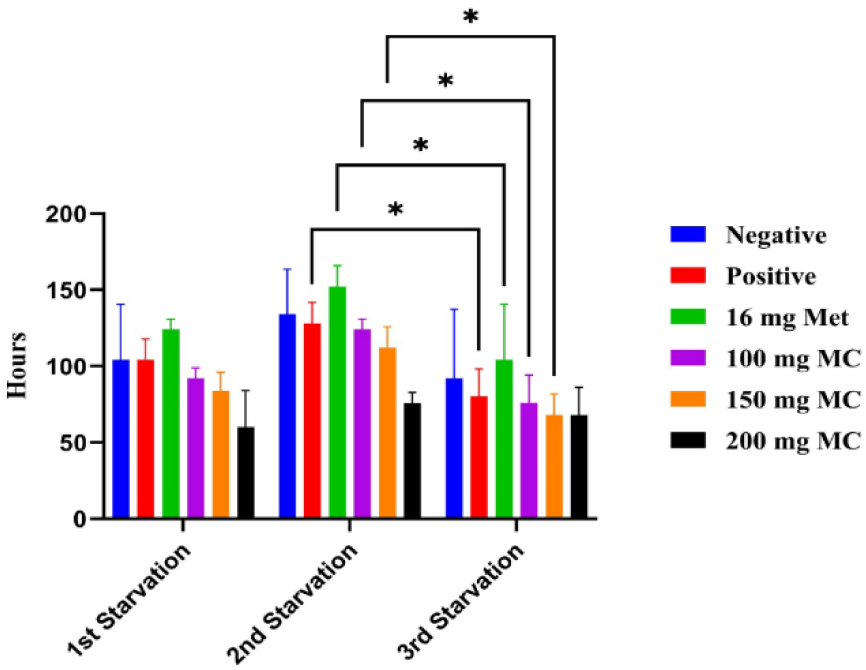
Male starvation resistance indigenous strains of *Drosophila melanogaster* induced with type 2 diabetes.

**Figure 1d:**
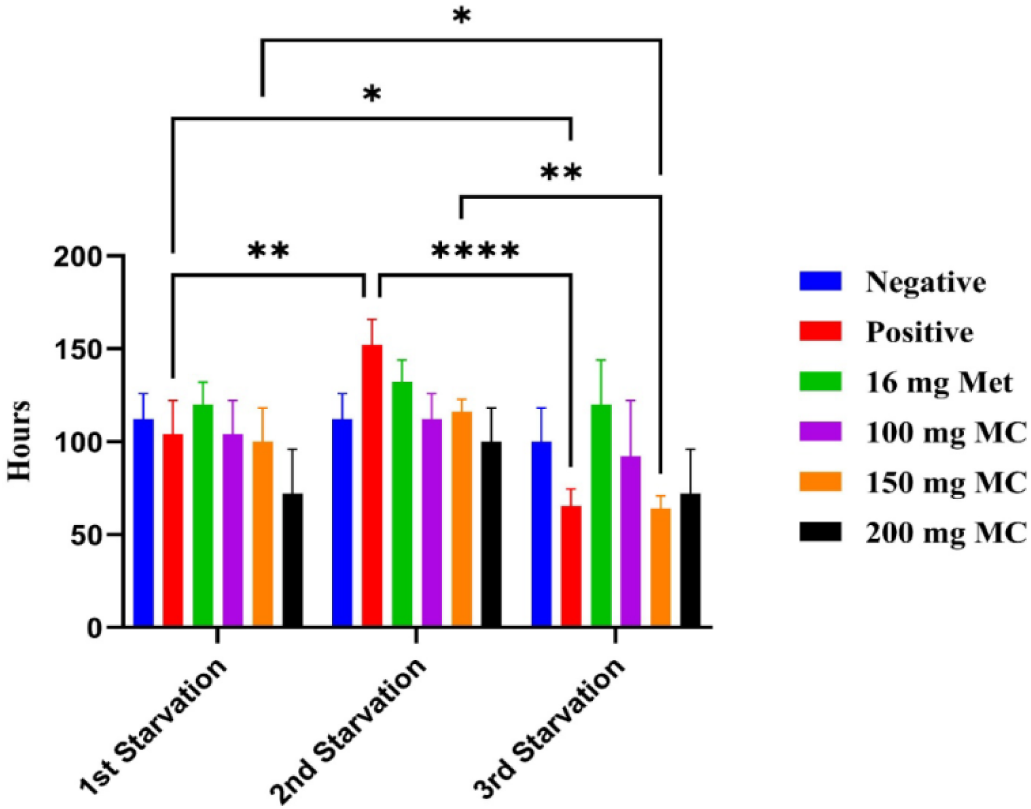
Females starvation resistance indigenous strains of *Drosophila melanogaster* induced with type 2 diabetes.

**Figure 1e:**
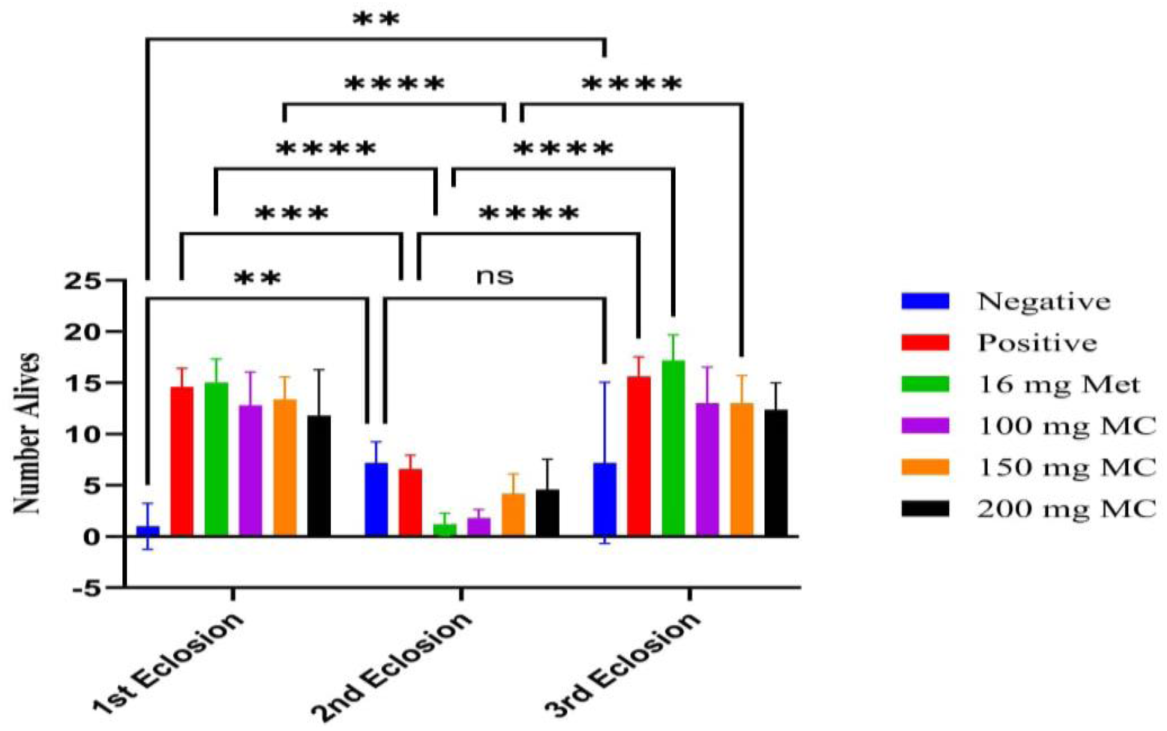
Eclosure rate of Harwich strain *Drosophila melanogaster* fed with metformin and *Mormodica charantia*.

**Figure 1f:**
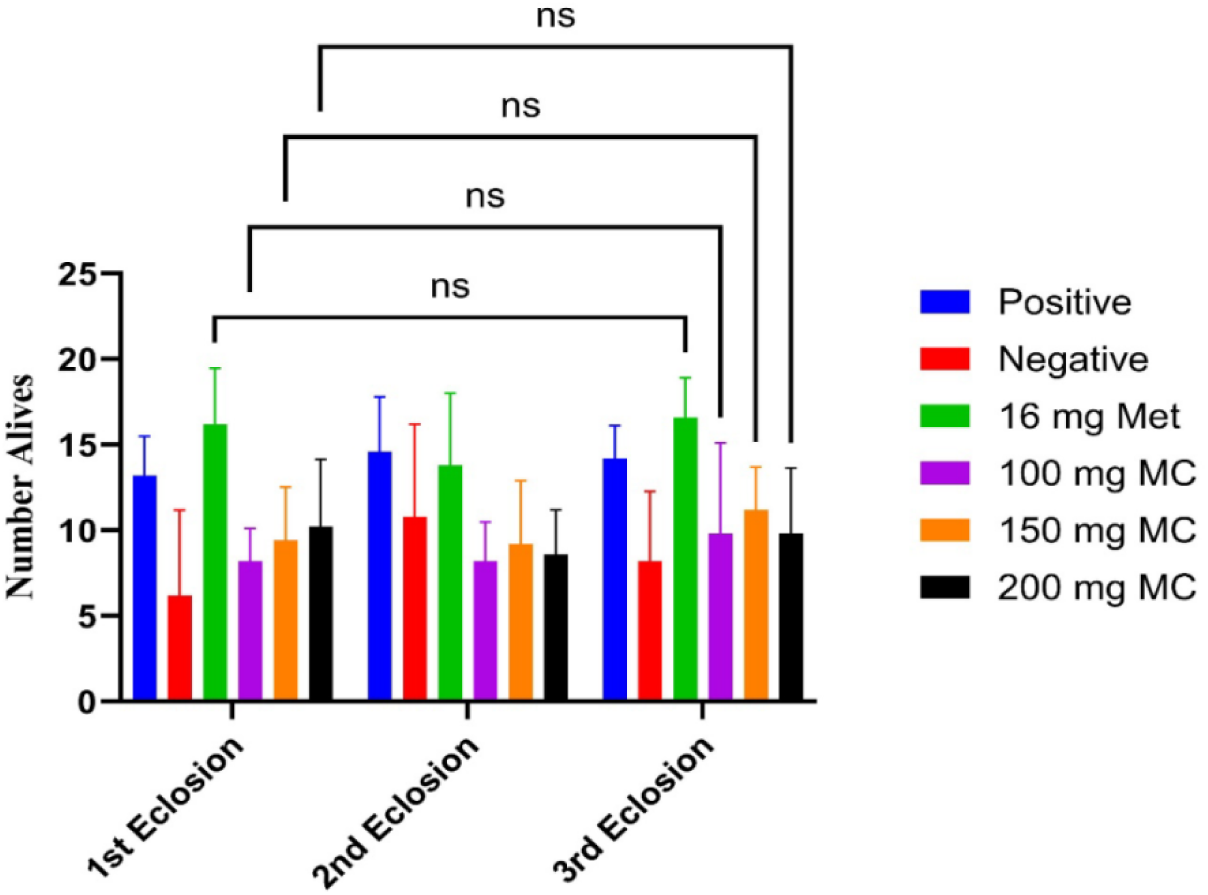
Eclosure rate of indigenous strain *D. melanogaster* induced with T2DM and fed with metformin and MC.

### Female Starvation resistance of Harwich strain *D. melanogaster* fed with metformin and *M. charantia*

The Negative and positive control group had the mean starvation resistance hours of (84 and 117.3) respectively, Metformin had the mean hours of flies that survived (94.7). Among the treatment group 100 mg had mean hours, survived for (102.7), 150 mg had (94.7) and 200 mg had mean hours survived for (92). The positive control group exhibited the highest starvation resistance, which survived for 92 and 117.3 hours, respectively. While the least survival rate was recorded in 100 and 200 mg for both male and female respectively (shown in figure 1b).

### Male Starvation Resistance of Diabetic indigenous *D. melanogaster* Treated With *M. charantia*

Figure 1c shows the male starvation resistance significant differences were observed between the control and treated groups. The negative control (non-diabetic untreated group) had the lowest starvation resistance across the three trials with a mean of approximately 60 hours, while the positive control (diabetic untreated group) had a higher starvation resistance, with a mean of approximately 70 hours. *M. charantia* at a dose of 0.15 g significantly improved (p < 0.05) the starvation resistance of the diabetic flies (75 hrs) when compared to other concentrations.

### Female Starvation Resistance of Diabetic indigenous *D. melanogaster* Treated With *M. charantia*

Figure 1d shows the female starvation resistance of *Drosophila melanogaster* induced with Type 2 Diabetes. Significant differences were observed between the control and treated groups. The negative control (non-diabetic untreated group) had the lowest starvation resistance across the three trials with a mean of approximately 55 hours, while the positive control (diabetic untreated group) had a higher starvation resistance, with a mean of approximately 65 hours. *M. charantia* at a dose of 0.15 g significantly improved (p < 0.05) the starvation resistance of the diabetic flies (mean of approximately 70 hours) when compared to other concentrations.

### Eclosure rate of Harwich strain *D. melanogaster* fed with metformin and *M. charantia*

The mean eclosion rate of T2DM-induced Harwich strain *D. melanogaster* was observed on the 11^th^ day post experimental setup, which is shown in (Figure 1e). Indicate that the negative and positive control group fed without MC had the mean number of eclosed flies (5.1 and 12.3) eclosed respectively; The 16 mg of Met had a mean number of eclosed flies (11.1). The 100 mg of MC had a mean number of eclosed flies (9.2) 150 mg had a mean number of another (10.2) eclosed, while 200 mg also had a mean number of eclosed flies (9.0). The highest mean number of eclosed flies was observed in the positive control group (12.3), while the least was observed in 200 mg of MC group (9.0). Among the treatment group 150 mg had the highest number of enclosed flies.

### Eclosion Rate of indigenous strain *D. melanogaster* Induced with T2D and Treated with *M. charantia*

Figure (1f) shows the eclosion rate of indigenous strain of *D. Melanogaster* induced with T2DM. Significant differences were observed between the control and treated groups. The negative control (non-diabetic untreated group) had the lowest number of eclosed flies across the three trials with a mean of 14 eclosed flies while the positive control (diabetic untreated group) had a higher eclosion rate, with a mean of 17 eclosed flies. *M. charantia* at a dose of 0.15 g improved (p > 0.05) the eclosion rate of the diabetic flies (mean of 17 eclosed flies) when compared to other concentrations.

#### Heat Tolerance

Heat tolerance assay showed no statistically significant differences in survival across treatments and strains (p > 0.05). In Harwich males, the highest survival (100%) was observed in the negative control, metformin (16 mg), and 200 mg *M. charantia*, while the lowest (66%) was recorded at 100 mg *M. charantia*. Among Harwich females, 100% survival was observed in the negative control and 150 mg *M. charantia*, while the lowest survival (60%) occurred with metformin and 200 mg *M. charantia* (Figure 2a).

**Figure 2a:**
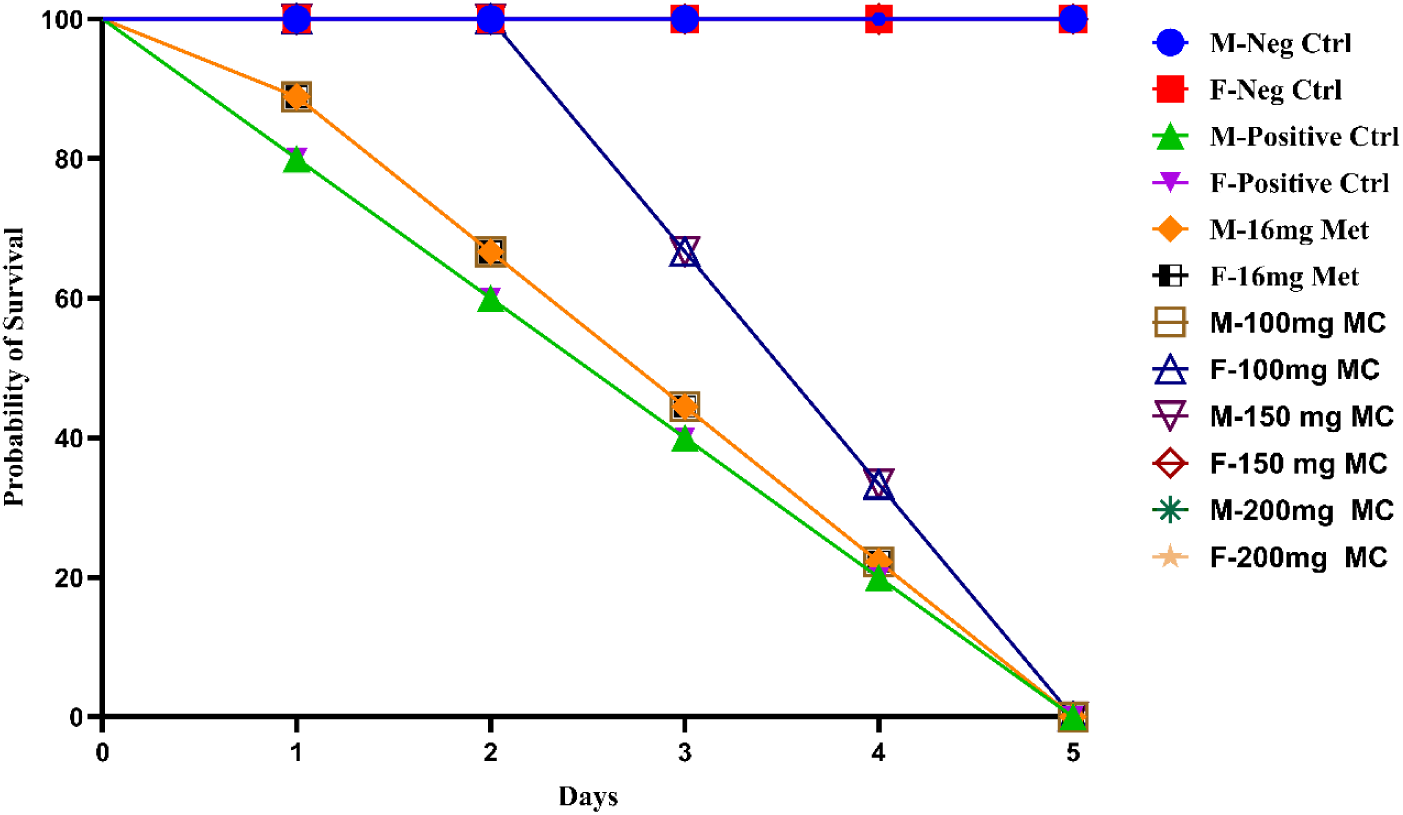
Heat tolerance survival curve of Harwich male and Female.

**Figure 2b:**
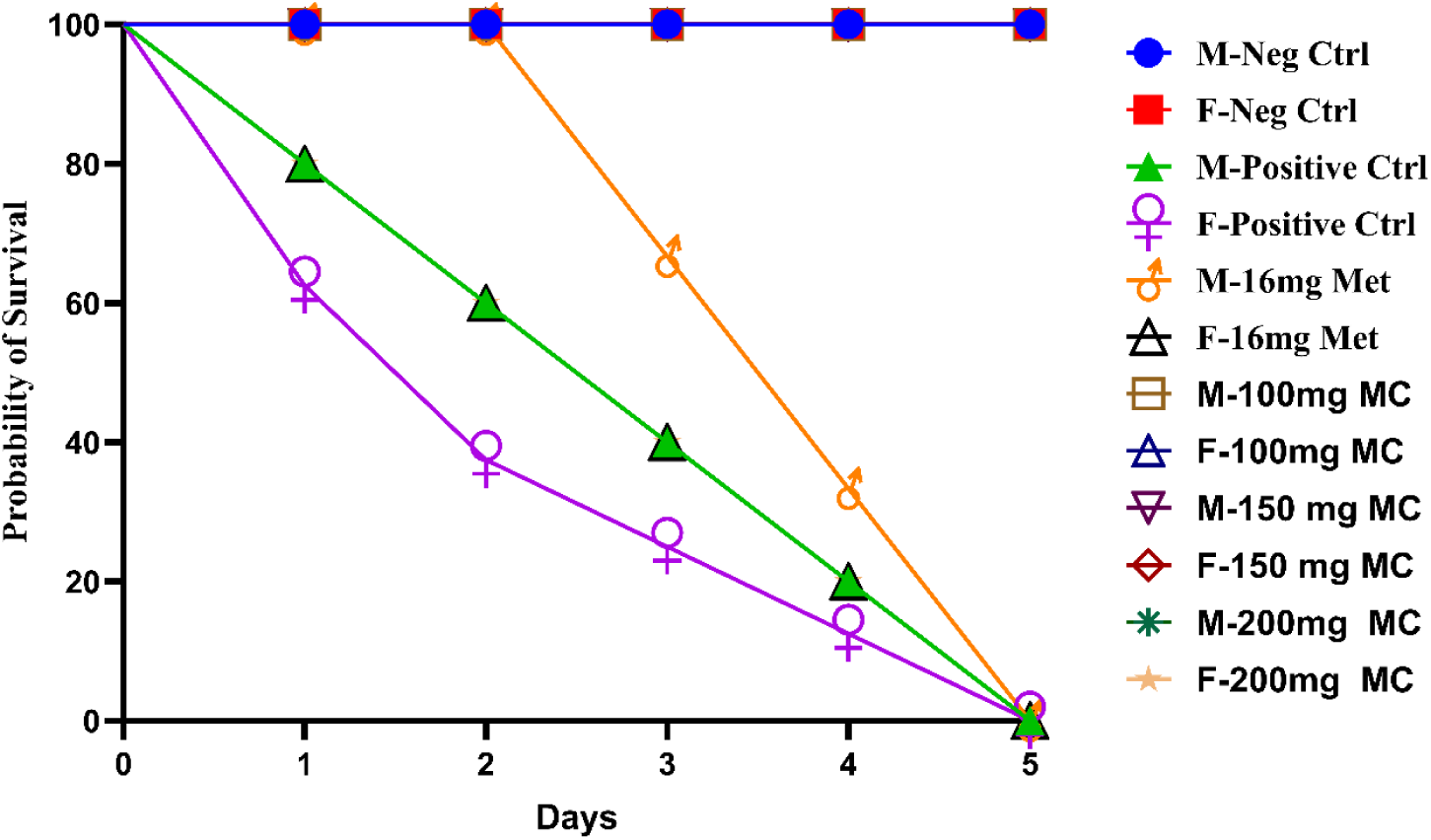
Heat tolerance survival curve of Ngd3 male and Female.

In Ngd3 males, the highest survival (100%) was seen in the negative control, 100 mg, and 150 mg *M. charantia*, with the lowest (90%) in the positive control, metformin, and 200 mg *M. charantia*. For Ngd3 females, the highest survival (100%) was recorded in the positive control and 150 mg *M. charantia*, while the lowest (60%) occurred in the negative control (Figure 2b).

### Effect of *M. charantia* on Malpighian tubule functions of Harwich strain *D. melanogaster* induced with type 2 diabetes

Figure 3(a) and (b) shows the effect of *M. charantia* on Malpighian tubule functions of Harwich strain *D. melanogaster* induced with type 2 diabetes. The positive control group shows increase in createnine level in both sexes (1.92 mg/dl and 1.59 mg/dl) when compared to the negative control (1.54 mg/dl) (1.33 mg/dl). Female Flies fed with 150 mg of MC has the lowest level of createnine (0.73 mg/dl) when compare among the groups. The reduction in Uric acid level was observe in male flies fed with 200 mg of MC (0.78 m/dl). Female flies show elevation in uric acid level in the positive group (1.06 m/dl). Sodium level in female flies, the positive control group shows an increase in sodium levels (1.67 m/dl), and among the treatment groups, the group fed 150 mg of MC has the lowest sodium levels (0.90 m/dl). Calcium level of both male (2.70mg/dl) and female (1.97 mg/dl) flies in the positive control group show increased in calcium levels. With those treated with 16 mg of metformin having the highest levels (1.68 mg/dl). However, among the treatment groups, 150 mg of MC has the lowest calcium levels in both male (3.67 mg/dl) and female (1.68 mg/dl) compare to the negative control group in both male (2.70 mg/dl) and female (1.98 mg/dl)

**Figure 3.**
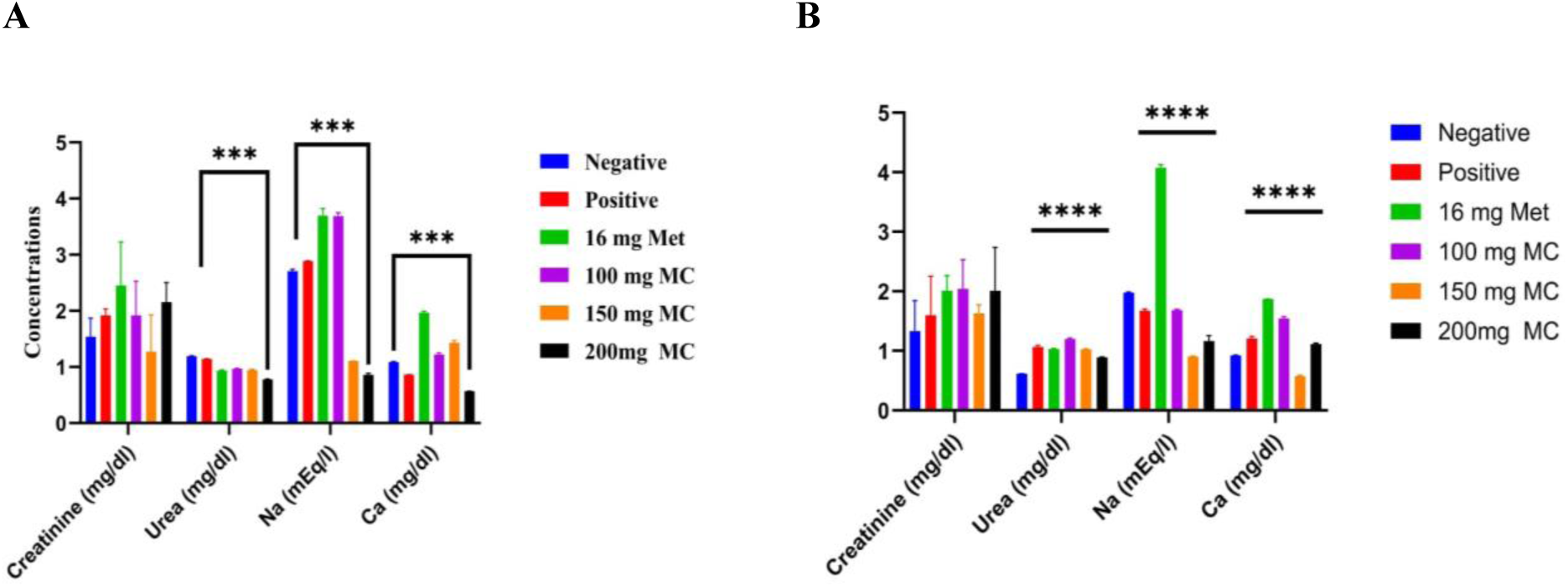
(a) and (b): Malpighian tubule function activity of Male and Female diabetic Harwich strain *D. melanogaster* induced with type 2 diabetes (Cr, Ur, Na and Ca)

**Figure: 3:**
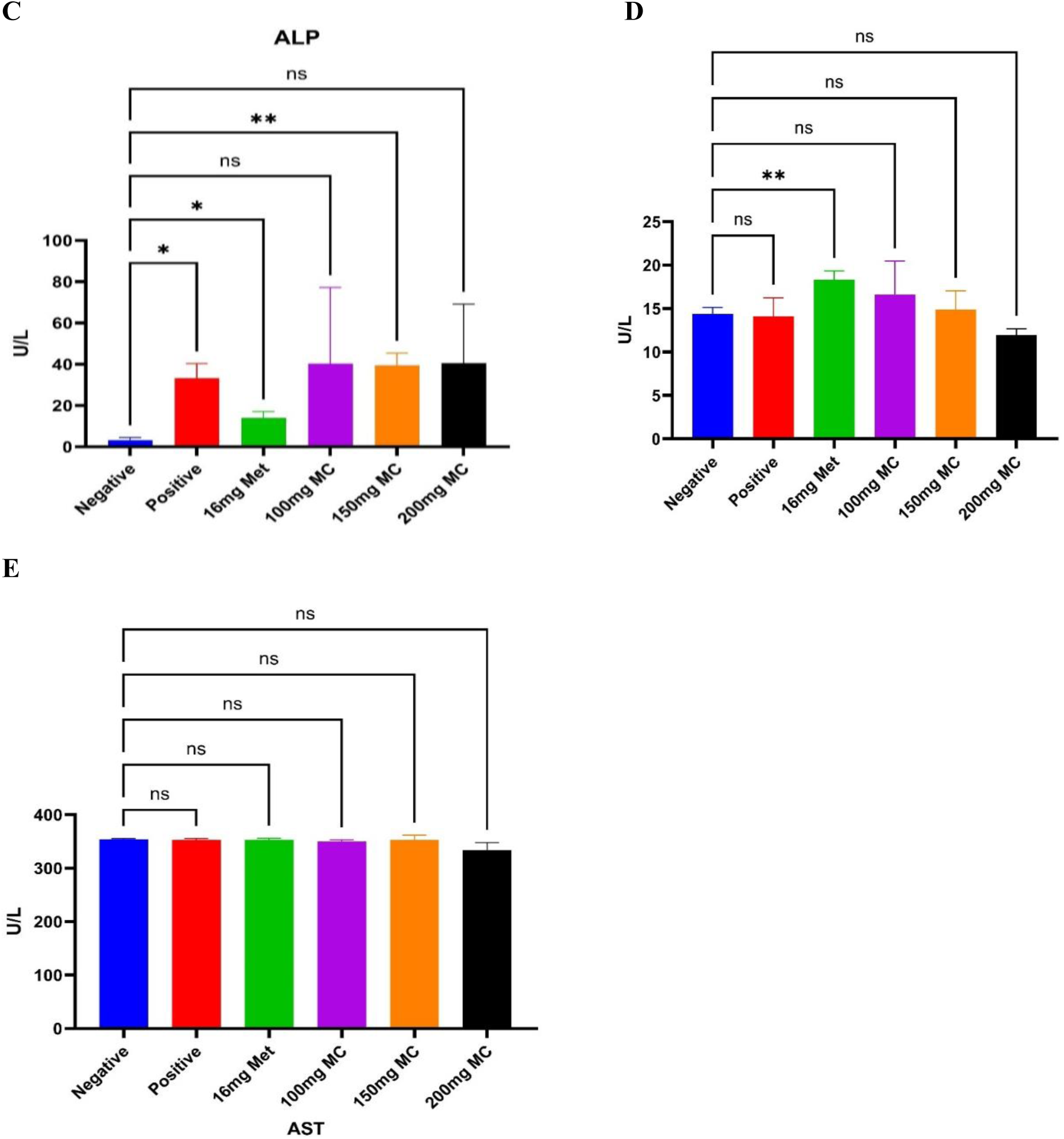
Fat body function of male diabetic Harwich Strain *D. melanogaster*, (C): Alkaline phosphate activity across the groups, (D) Alanine aminotransferase acidity across the groups, (E): Aspartate aminotransferase activity across the groups.

**Figure: 3.**
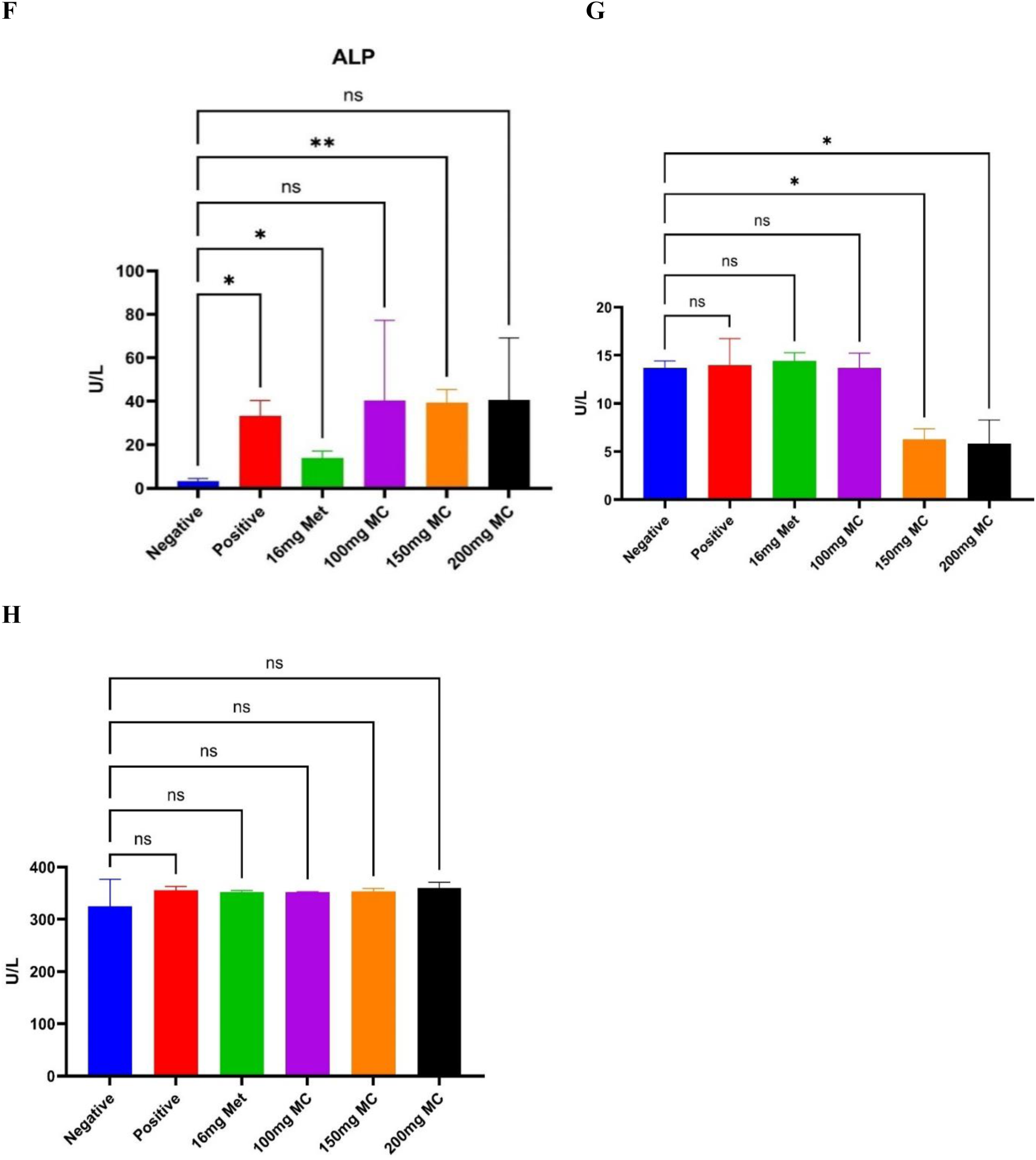
Fat body function of female diabetic Harwich Strain *D. melanogaster*, F: Alkaline Phosphatase activity across the groups, G: Alanine aminotransferase acidity across the groups, H: Aspartate aminotransferase activity across the groups.

### Effect of *M. charantia* on fat body function of Harwich strain *Drosophila melanogaster* induced with type 2 diabetes

In Figures 3(c) and 3(d), both male (3.33 µ/L) and female (33.33 µ/L) flies shows increase in ALP level in positive group with flies fed with 100 mg of MC having the highest ALP level male (9.09 µ/L) and female (40.30 µ/L). Negative group flies have the lowest ALP level male (19.09 µ/L) and female (3.33 µ/L). Figures 3(e) and 3(f) shows similar level of ALT in the positive group male (14.61 µ/L) and female (13.97 µ/L) and negative group in both male (14.00 µ/L) and female (13.68 µ/L). With 16 mg of MET having the highest level (18.33 µ/L) in male flies. Flies fed with 200 mg of MC shows the least level (14.04 µ/L) of ALT in male. There was significant reduction in female flies fed with 150 mg of MC (6.27 µ/L) and 200 mg of MC (9.89 µ/L). Figures 3(g) and 3(h) showing similar level of AST in both the control group and treatment group of male flies with 200 mg of MC having the lowest level (334. u/l) and negative group having the highest level (354 µ/L). Evident increase show in positive group of female flies (355.6 µ/L), with 200 mg of MC having the highest level (360.3 µ/L).

### Effect of *M. charantia* on Oxidative Stress function of Harwich strain *Drosophila melanogaster* induced with type 2 diabetes

Oxidative Stress Function Activity, Figure 4 (a) and (b). There was decrease in MDA level in female flies of the positive group (2.01 µ/ml). Male flies fed with 200 mg of MC has the highest level (3.94 µ/ml) of MDA. In flies fed with 100 mg of MC (3.18 µ/ml) having the least value among the MC treatment groups of female. Positive group flies have the highest CAT level in male flies (5.33 u/ml), with 200 mg of MC, having the least level of CAT in male (2.64 µ/ml). There was significant increase in CAT level in negative group of female flies (5.88 µ/ml) when compared to positive group (4.94 µ/ml),

**Figure: 4.**
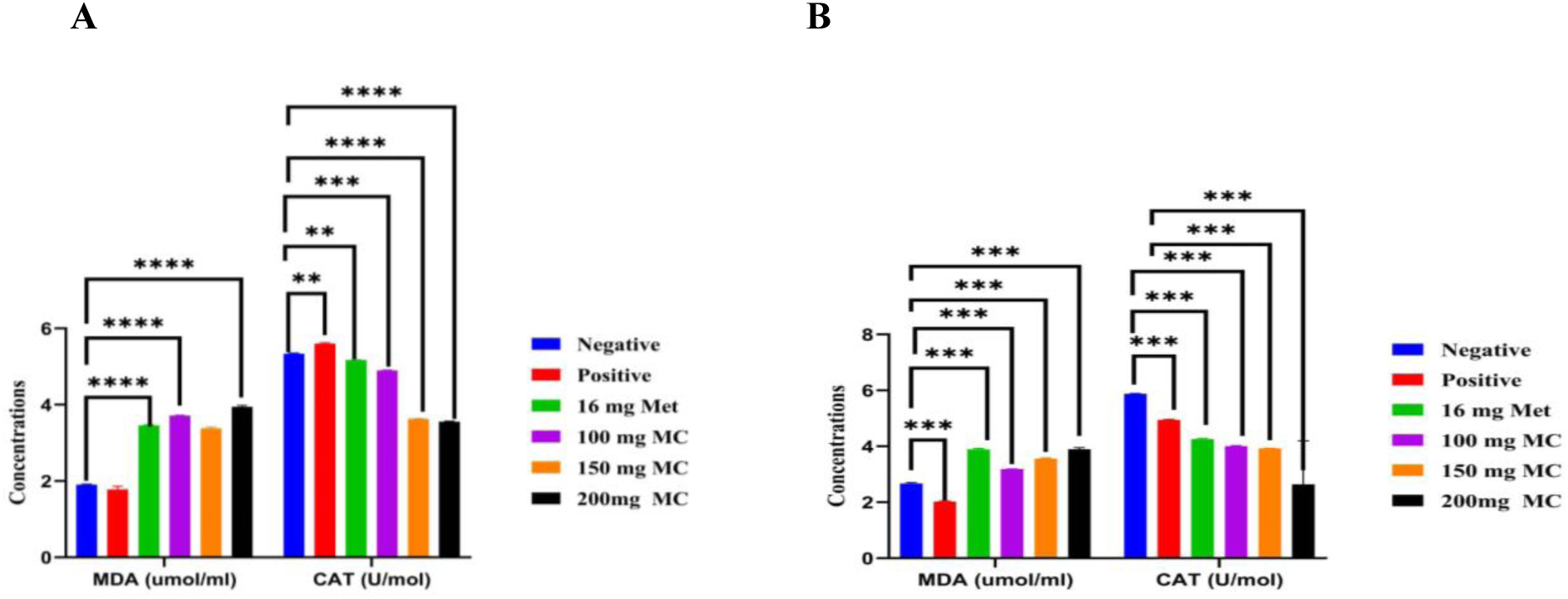
(a) and (b). Oxidative stress of Male and Female Harwich strain *D. melanogaster* induced with type 2 diabetes. (MDA and CAT)

**Figure: 4a.**
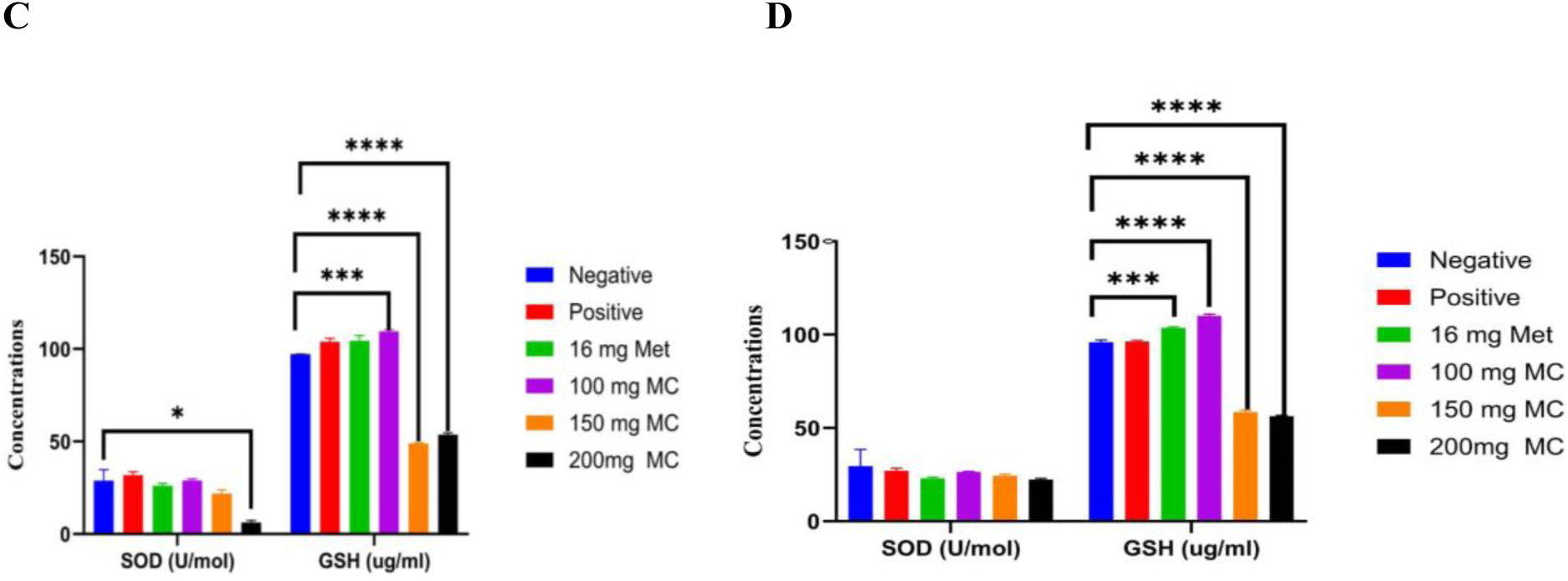
(C) and (D): Oxidative stress of Male and female Harwich strain *D. melanogaster* induced with type 2 diabetes. (SOD and GSH)

Figure 4 (c) and (d) shows Positive group shows evident increase in SOD level (28.48 µ/ml) in male flies, with flies fed with 200 mg of MC having the least SOD level (6.1 µ/ml). There was decrease in SOD level in female flies of the positive group (27.1 µ/ml) compared to negative group (29.53 µ/ml) with 100 mg of MC showing a higher level (26.90 µ/ml) of SOD among the treatment groups of female. Male positive group flies shows evident increase in GSH level (97.16 µ/ml) when compared to the negative group (95.95 µ/ml), with 100 mg of MC having the highest GSH level in male flies (109.46 µ/ml). 150 mg of flies having the least GSH level among the groups (48.77 µ/ml). Significant increase was observed in female flies fed with 100 mg of MC (101.02 µ/ml), when compared to positive control group (96.29 µ/ml).

### Transcriptional Regulation of the *CG4607* gene in Response to *Momordica charantia* Treatment

Figure (5a) shows the relative fold changes in *CG4607* gene expression across treatment groups in both Harwich and indigenous strains of *D. melanogaster.* In the Harwich strain, the highest upregulation of *CG4607* was observed at 200 mg *M. charantia* (-0.5665), while the highest downregulation occurred at 150 mg *M. charantia* (1.621). In the Ngd3 strain, the highest upregulation was seen in the positive control (-1.543), whereas the highest downregulation was recorded at 150 mg *M. charantia* (5.777).

**Figure: 5a.**
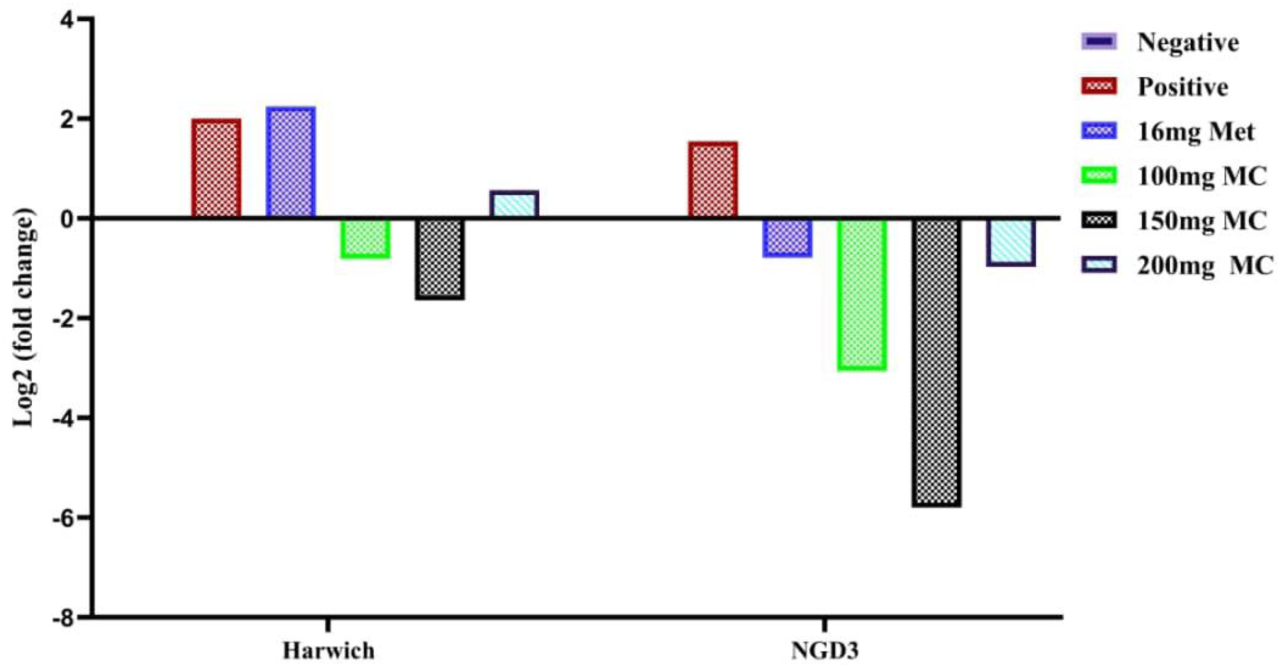
*CG4607* Gene Expression level in Harwich and indigenous (Ngd3) N: Negative control, P: Positive control, T: Treatment.

**Figure: 5.**
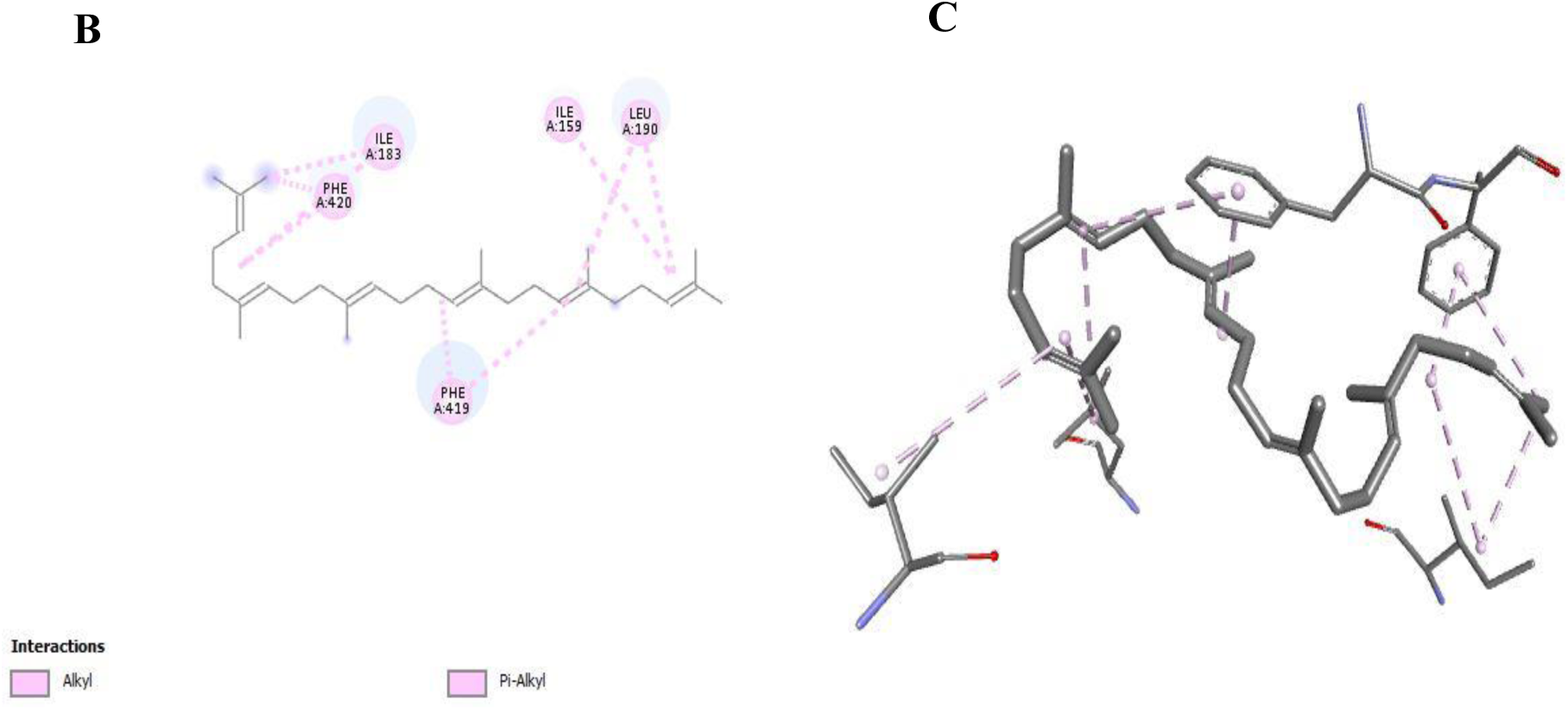
The (B) 2D and (C) 3D interaction between squalene and *CG4607 gene* modelled protein structure.

**Figure 5:**
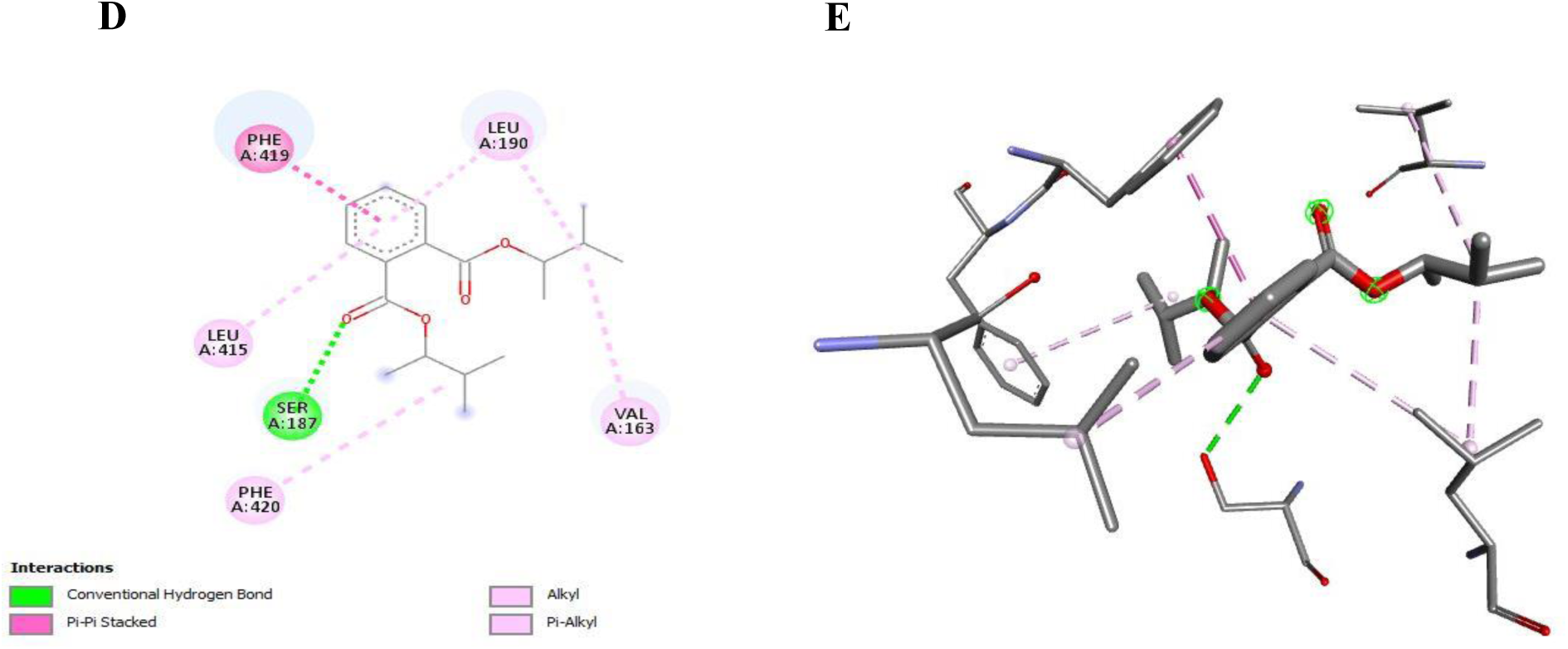
The (D) 2D and (E) 3D interaction between Bis (3-methylbutan-2-yl) phthalate and *CG4607* modelled protein structure.

### Molecular docking of *Momordica charantia* compounds to the *CG4607* gene

The Ramachandran plot (Figure 4.1) confirmed a high-quality CG4607 protein model with 94.0% of residues in allowed regions. Among the docked compounds, Squalene showed the highest binding affinity to *CG4607* (-7.5 kcal/mol), forming eight hydrophobic interactions with Phenylalanine, Leucine, and Isoleucine residues and Bis(3-methylbutan-2-yl) phthalate showed higher binding affinity to *CG4607* (-7.4 kcal/mol) (Figure 4.2a and Figure 4.2b). The lowest binding affinity was observed with 4-Fluoro-1-methyl-5-carboxylic acid, ethyl ester (-4.5 kcal/mol) (Table 1:).

**Table: 1.** Molecular Docking of *M. Charantia* Compounds Using *Drosophila* Glucose transporter (*CG4607*)

| S/N | Compounds | Binding Energies (kcal/mol) |
| --- | --- | --- |
| 1 | 10-Undecenal | -4.6 |
| 2 | Oxirane, [(dodecyloxy)methyl]- | -4.6 |
| 3 | 2-Piperidinone, N-[4-bromo-n-butyl]- | -5.0 |
| 4 | 9,12-Octadecadienal | -5.3 |
| 5 | Ethyl 13-methyl-tetradecanoate | -5.4 |
| 6 | **Bis(3-methylbutan-2-yl) phthalate** | **-7.4** |
| 7 | **Squalene** | **-7.5** |
| 8 | 4-Fluoro-1-methyl-5-carboxylic acid, ethyl(ester) | -4.5 |
| 9 | Z-8-Pentadecen-1-ol acetate | -5.4 |
| 10 | Cyclohexanol, 2-(2-propynyloxy)-, trans- | -4.9 |
| 11 | 1-Methoxymethoxy-oct-2-yne | -4.6 |
| 12 | 2-Methyl-Z,Z-3,13-octadecadienol | -5.3 |
| 13 | 7,11-Hexadecadienal | -5.9 |
| 14 | cis-Vaccenic acid | -5.6 |
| 15 | 9-Oxabicyclo[6.1.0]nonane, cis- | -4.8 |
| 16 | Dispalure | -5.6 |
| 17 | Hexadecanoic acid, ethyl ester | -6.1 |
| 18 | Oleic acid | -5.4 |
| 19 | trans-Traumatic acid | -5.1 |
| 20 | Undec-10-ynoic acid, tetradecyl ester | -5.7 |
| 21 | Z-17-Nonadecen-1-ol acetate | -6.4 |
| 22 | Oxalic acid, cyclobutyl hexadecyl ester | -6.3 |

#### Electropherograms

The amplified bands of *CG4607* gene in Harwich Strain *D. melanogaster* is shown in Plate II, and plate III shows the Electropherogram of indigenous Strain *D. melanogaster* their band sizes are 138bp respectively.

#### Genetic Variations and Phylogenetic Relationships of the CG4607 gene Among T2DM-induced *Drosophila melanogaster* Strains treated with *Momordica charantia*

Amplified bands of 138 bp were observed in both strains (Plates II-III). Based on multiple sequence alignment (MSA), the negative controls (HN, NN) displayed conserved sequences with 0 mutations, while the diabetic controls (HP, NP) exhibited multiple sequence variations, with 7 mutations in HP and 9 in NP. Metformin-treated groups (H16, N16) showed fewer sequence variations than the diabetic controls, with H16 showing 3 mutations and N16 showing 4 mutations. *M. charantia* treatments induced dose-dependent variations. The lowest number of mutations was recorded in H100 (1) and N100 (2), while the highest number of mutations was observed in H200 (6) and N200 (8). Ngd3 strain groups (N100–N200) showed more sequence alterations overall compared to Harwich groups (H100–H200) as shown in Figure (6a).

Figure (6b) presents the phylogenetic tree generated from *CG4607* gene sequences. HN and NN clustered separately, while HP and NP grouped closely. Metformin-treated groups (H16, N16) appeared as distinct branches. Among *M. charantia*-treated groups, N100–N200 formed a single cluster (bootstrap: 59%), and H100–H150 clustered together (bootstrap: 54%). H200 and N200 formed independent branches with bootstrap values of as shown in (61%) and (66%), respectively.

**Figure 6a:**
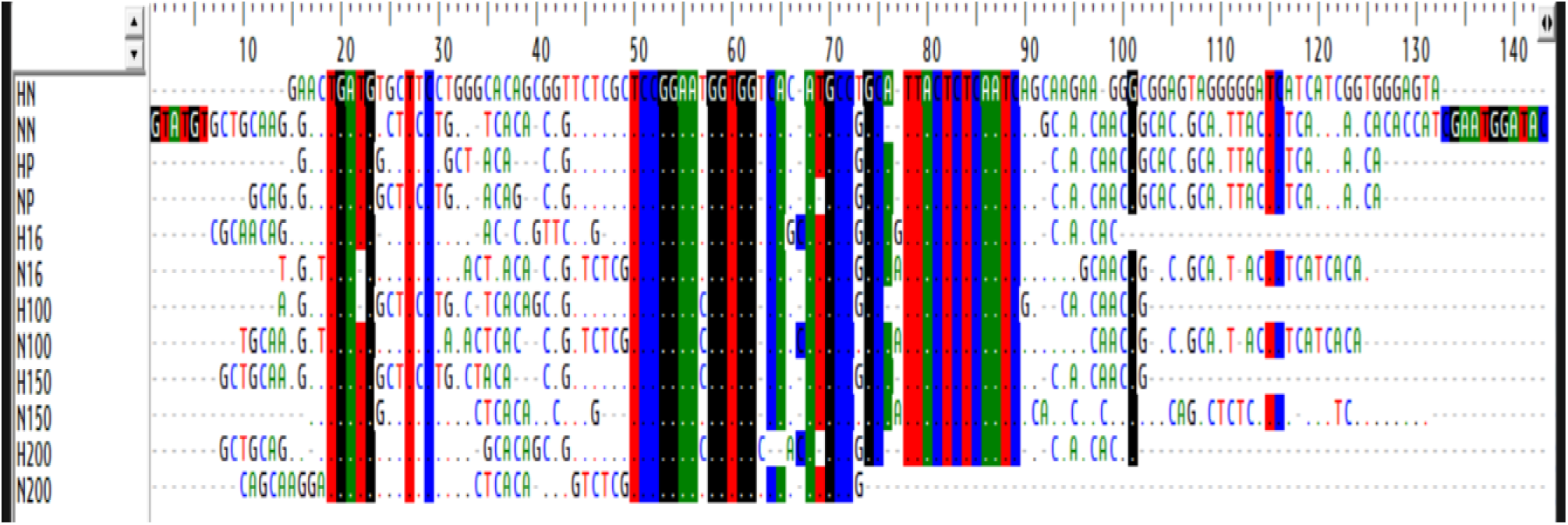
Multiple Sequence Alignment (MSA) of Glucose Transporter Gene (CG4607) of Harwich and Ngd3 strains of *D. melanogaster* induced with type 2 diabetes and treated with *M. charantia*.

**Figure 6b:**
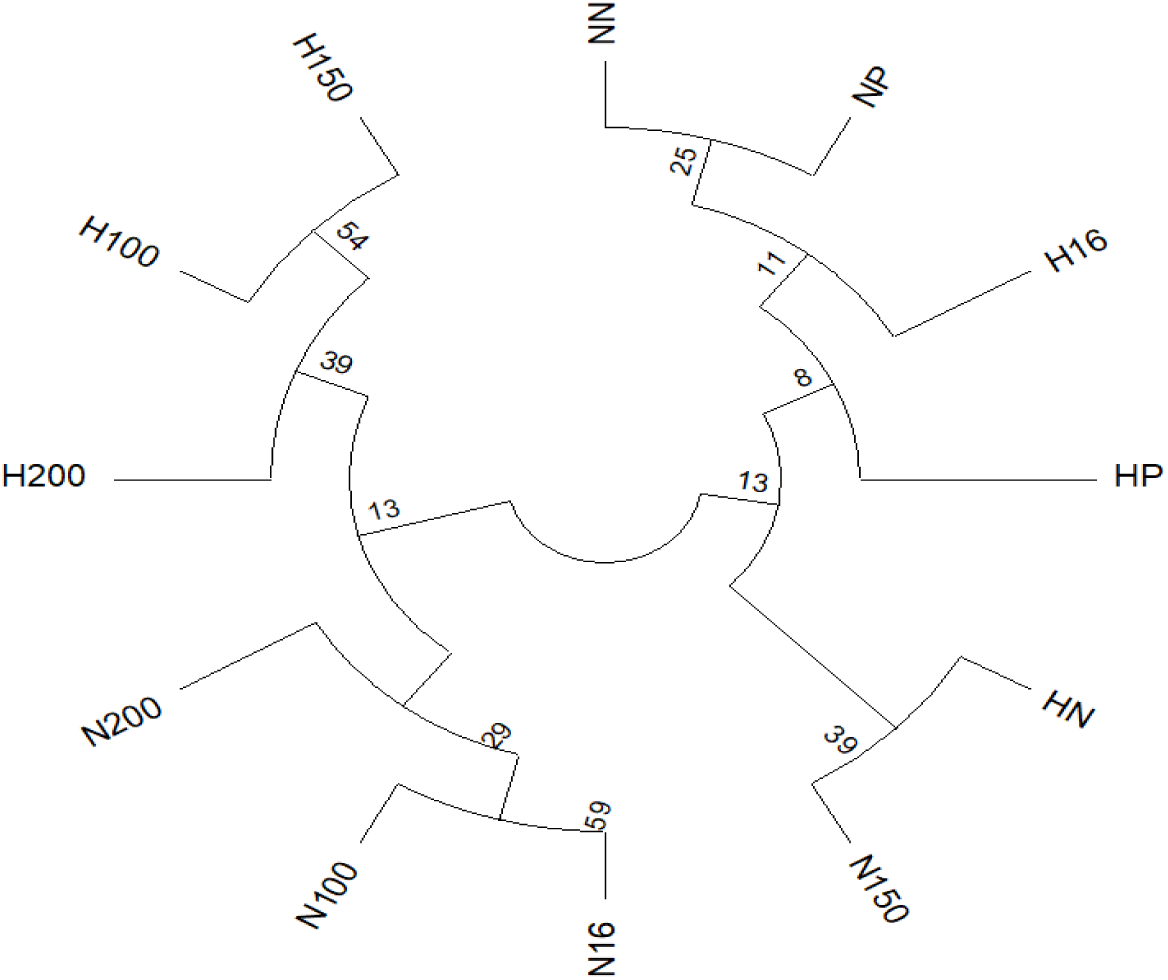
Phylogenetic tree of Genetic variation of Harwich and Ngd3 strains of *D. melanogaster* induced with type 2 diabetes and treated with *M. charantia*. N: Ngd3, H: Harwich, HN: Harwich negative.

**Plate II:**
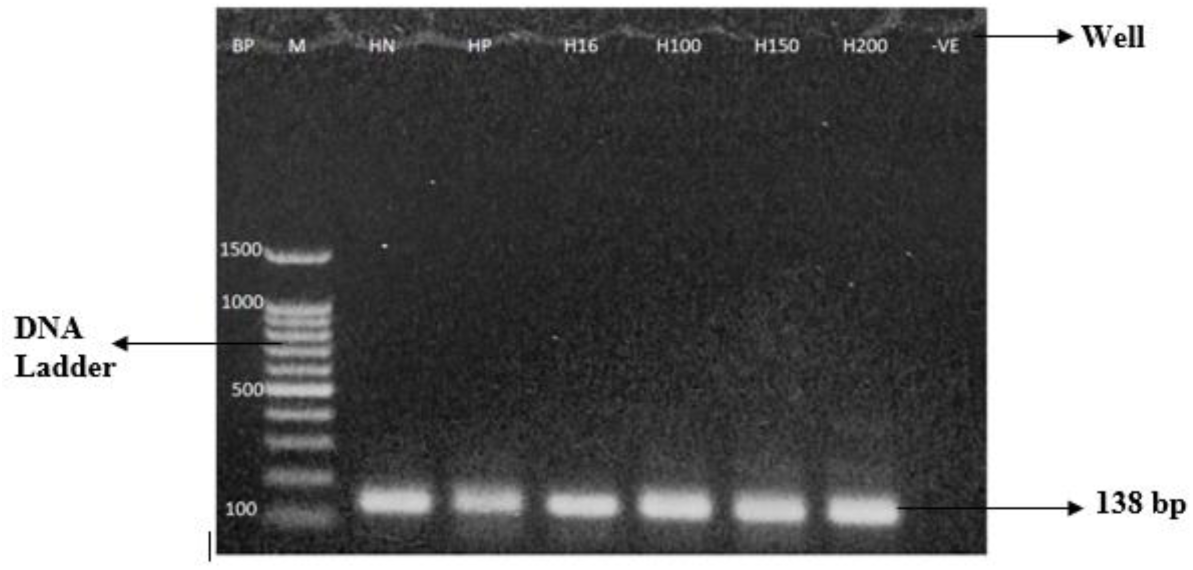
Electropherogram of *CG4607* Gene of Harwich Strain *Drosophila melanogaster*. BP: Base pair, M: DNA Ladder 100bp, HN: Harwich Negative control, HP: Harwich Positive control, H16: Harwich 16 mg Metformin, H100: Harwich 100 mg, H150: Harwich 150 mg, H200: Harwich 200 mg.

**Plate III:**
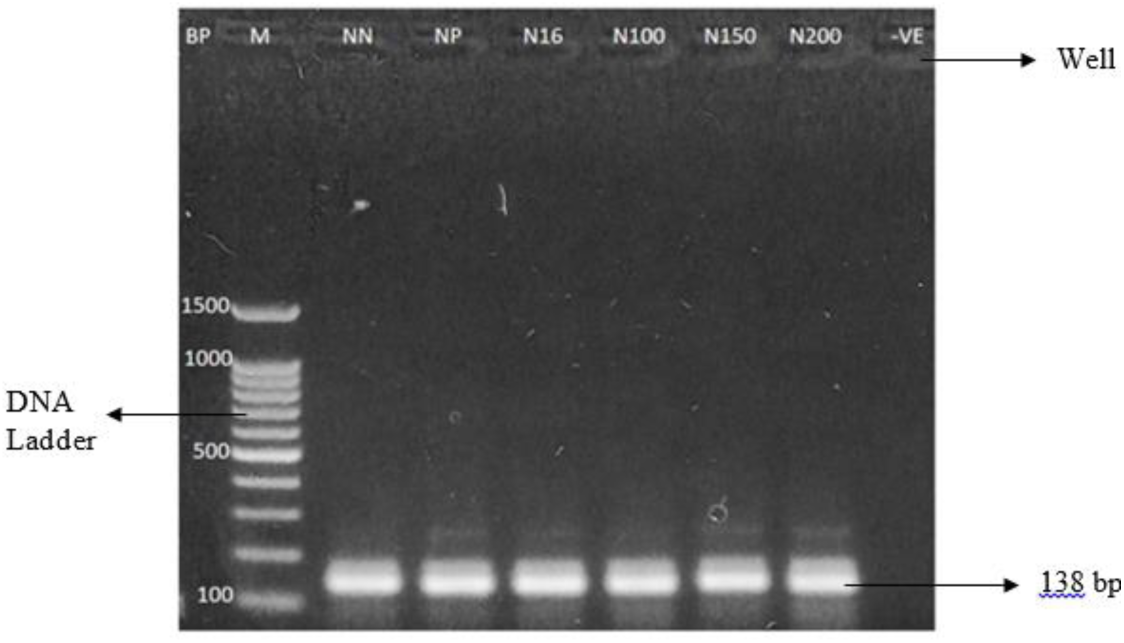
Electropherogram of *CG4607* Gene of Ngd3 Strain *Drosophila melan aster*. BP: Base pair, M: DNA Ladder 100bp, NN: Ngd3 Negative control, NP: Ngd3 Positive control, N16: Ngd3 16 mg Metformin, N100: Ngd3 100 mg, H150: Ngd3 150 mg, N200: Harwich 200 mg.

## DISCUSSION

The control and treatment groups levels of hunger resistance clearly differed, this study shows Harwich strain diabetic flies, the positive control group showed the greatest resilience to starvation. Which implies that T2DM induced metabolic alterations, could be due to of changed fat storage and metabolic pathways, provide a certain level of resistance to hunger. Diabetic flies frequently show enhanced lipid storage, which may improve their capacity to endure starvation (Musselman *et al*., 2011), insulin resistance and dysregulated glucose metabolism in type 2 diabetes cause energy to be stored as fat, which extends survival periods during nutritional shortage. Flies fed with metformin showed better resistance to starvation than the negative control, but not as much as the positive control. Metformin assisted in regulating energy metabolism in diabetic flies, improving glucose metabolism and reducing fat accumulation, Metformin-treated flies might have reduced lipid stores compared to untreated diabetic flies, thus showing lower starvation resistance. This observation is consistent with findings by Foretz *et al*. (2014), who explored Metformin’s metabolic effects.

In Harwich strain starvation resistance Males (89.3 hours) and females (102.7 hours) exhibited the strongest hunger resistance among the treatment groups when given 150 mg and 100 mg of *M. charantia*. The survival times were shorter at 200 mg, though, with the male group exhibiting the least amount of starvation resistance (69.3 hours) and the female group exhibiting the least amount (92 hours), this impact is dose-dependent and consistent with previous studies on MC. It has been explored that moderate dosages of MC control lipid and glucose metabolism, which may improve energy consumption during starvation. According to research by Joseph and Jini (2013), *M. charantia* can improve metabolic flexibility by affecting the mobilization and storage of lipids. At higher dosages (200 mg), some substances, such as alkaloids and saponins, may have toxic effects that interfere with regular metabolic processes, lowering resistance to starvation. The findings also show that starvation resistance varies between sex, with females in all groups often outliving males. This might be explained by variations in the sexes’ energy needs, fat storage, and metabolic rates. According to research, female *Drosophila* have a tendency to store more fat than males, which may help them survive longer periods of starvation. Wu *et al*. (2017) discovered that because of their larger bodies and energy stores, females *Drosophila* frequently exhibit higher starvation resistance.

In indigenous strains the comparison of starvation resistance between the diabetic untreated group and the non-diabetic untreated group reveals that the positive control consistently showed higher starvation resistance in both male and female *D. melanogaster*. This suggests that the diabetic condition may have prompted metabolic adaptations that enhanced survival under starvation conditions. The positive response in the MC treated group at a dose of 150 mg further supports the hypothesis that MC can alleviate the adverse effects of diabetes-like conditions, maintaining higher starvation resistance similar to the positive control group for both sexes. For male flies, the 100 mg of MC dose showed generally lower starvation resistance compared to the 150 mg group but was still better than the negative control, indicating mild survival benefits at this lower dose. However, the 200 mg dose resulted in lower starvation resistance across all groups, suggesting a dose-dependent effect. This pattern is consistent with findings from Miao *et al*. (2022), which reported variable outcomes with different dosages of natural extracts, highlighting the importance of optimizing dose for therapeutic effectiveness. Similarly, for female flies, the 100 mg dose showed generally lower starvation resistance compared to the 150 mg group but was still better than the negative control, indicating mild survival benefits at this dose. However, the 200 mg dose resulted in lower starvation resistance across all groups, suggesting a dose-dependent effect, which aligns with Zhang *et al*. (2023) who found that different dosages of natural extracts can lead to varying physiological responses. Their study emphasized the need for precise dosage to achieve optimal therapeutic benefits.

The eclosion rate was significantly impacted by the induction of T2DM in Harwich strain *D. melanogaster*. The significant eclosion rate was seen in the positive control group, which was fed without MC, and reduced eclosion was seen in the non-diabetic flies. The 150 mg group had the highest eclosion rate among the treatment groups, indicating a dose-dependent response to MC treatment. However, the eclosion rate was lowest in the 200 mg group; this implies that whereas low dosages of MC may support developmental health, greater dosages may have an inhibitory effect. Compounds such as polypeptide-p, vicine, and charantin found in MC are known to have hypoglycemic effects at modest dosages. However, Joseph and Jini (2013) showed that large levels of MC can cause oxidative stress, which can impair cellular functioning, suggesting that high doses can be hazardous. Also, Grover and Yadav (2004) emphasize that while MC has hypoglycemic effects at therapeutic doses, excessive intake can lead to adverse side effects, which corroborates the reduced eclosion rates seen at the 200 mg dose.

The heat tolerance assay, a standard survival analysis in *D. melanogaster* research used in assessing the effects of heat stress on the lifespan of flies, and the underlying genetic, molecular, and physiological mechanisms of heat tolerance (Sørensen *et al*., 2003; Le Bourg and Moreau, 2014). This assay, revealed potential protective effects of *M. charantia* against thermal stress. Groups treated with 100 mg and 150 mg of the extract showed 100% survival, outperforming both the positive control and Metformin-treated groups, which experienced some mortality. Although differences among treatment groups were not statistically significant, these findings suggest that *M. charantia* may confer modest thermal protection, potentially through antioxidant mechanisms. A sex-specific survival trend was also observed, with male Ngd3 flies exhibiting greater resilience, warranting further investigation into sex-linked stress responses. Several studies have highlighted the role of *M. charantia* in enhancing stress resistance, including thermal stress. For instance, a study by Kumar *et al*. (2013), observed that extracts of *M. charantia* improved the thermal tolerance of diabetic rats by modulating the antioxidant defense system, reducing oxidative stress, and maintaining cellular homeostasis. A sex-specific survival trend was also observed, with male Ngd3 flies exhibiting greater resilience. Although differences among treatment groups were not statistically significant, these findings suggest that *M. charantia* may confer modest thermal protection, potentially through antioxidant mechanisms. This could imply that the basal stress resistance in *D. melanogaster* is already high, or that the protective effects of *M. charantia* are only marginally better than the normal physiological response to heat stress (Vesa *et al*., 2024).

The creatinine levels in the positive control group showed an increase with aligns with findings from Patil *et al*., (2023), who described that oxidative stress leads to renal dysfunction in *D. melanogaster*, as indicated by elevated creatinine levels. In contrast, the reduced creatinine levels observed in female flies treated with 150 mg of MC suggest a protective effect, potentially attributed to the antioxidant properties of MC. Which could indicated that MC can mitigate kidney function in metabolic disorders (Patil *et al*., 2023). The reduction of uric acid in male flies treated with 200 mg of MC corresponds with research by Kumar *et al*. (2024), which noted that certain dietary interventions could mitigate hyperuricemia in model organisms. Elevated uric acid levels in female positive control flies reinforce the connection between oxidative stress and metabolic dysregulation, as discussed by Gao *et al*. (2022), stated that oxidative stress is a significant contributor to elevated uric acid levels in diabetic models. The observed decrease in sodium levels in female flies treated with 150 mg of MC is consistent with studies showing that antioxidants can restore ion homeostasis disrupted by oxidative stress (Zhao *et al*., 2020). Thus, MC may play a role in maintaining sodium balance during oxidative stress, corroborating findings from a recent study by Zhang *et al*. (2022) that reported similar effects of plant extracts on ion regulation in *D. melanogaster*. The increase in calcium levels in the positive control group and the modulation seen with MC treatment supports the work of Ahmed *et al*. (2022), found out that calcium homeostasis is disrupted during oxidative stress in *D. melanogaster*. The significant reduction in calcium levels in the 150 mg MC treatment group highlights MC potential role in restoring calcium balance, as noted by Rehman *et al*. (2023), where found that plant extracts could ameliorate calcium dysregulation in metabolic models.

Sodium level decreases in flies fed 150 mg and 200 mg of MC suggests that MC helps in balancing electrolyte in diabetic conditions; the increase in calcium levels in the treatment groups indicates that MC was able to suppress high, and supporting Malpighian tubule function. Singh *et al*., (2021) observed reduction in ALP levels in MC treated groups, particularly with 100 mg, suggests a hepatoprotective effect. A study by Senanayake *et al*. (2004) showing that MC extract reduced liver enzyme levels in oxidative stress models, consistent with its anti-inflammatory properties. ALT reduction in females treated with 150 mg and 200 mg MC was as a result of hepatic antioxidant defence which aligns with the work of Abdulazeez *et al*. (2024), found that MC lowers ALT levels by enhancing hepatic antioxidant defenses. This suggests MC could have potential to mitigate liver damage through antioxidant action. The AST increase in females flies fed with 200 mg MC can be linked to sex-specific metabolic responses, as supported by Hamden *et al*., (2008), highlighted that oxidative stress induces different biochemical reactions in male and female *D. melanogaster*. The decrease in MDA levels in female flies of the positive group suggests a reduction in lipid peroxidation, a common indicator of oxidative stress. also with research by Gohil *e*t *al*. (2022), reported that antioxidant treatments can significantly reduce MDA levels, indicating a protective effect against oxidative damage. Conversely, the significant increase in MDA levels in male flies fed with 200 mg of MC indicates a potential effect at higher doses, which has been documented in previous studies where high concentrations of phytochemicals led to increased oxidative stress (Mansoor *et al*., 2023). Female flies fed with 100 mg of MC exhibited the lowest MDA levels among the treatment groups, reinforcing the idea of a dose-dependent response. The observed decrease in CAT levels in the positive control group males, particularly those treated with 200 mg of MC could be an impairment in the antioxidant defense mechanism, which aligns with findings by Zhang *et al*. (2022), noted that excessive oxidative stress can overwhelm antioxidant enzymes, reducing their activity. In contrast, the significant increase in CAT levels in the negative group of female flies indicates an adaptive response to maintain oxidative balance. This observation is critical, as maintaining CAT levels is essential for cellular protection against oxidative damage (Agarwal *et al*., 2021).

The evident increase in SOD levels in the positive group male flies aligns with the findings of Boonchird *et al*. (2023), where oxidative stress led to the upregulation of SOD as a compensatory mechanism. However, the reduced SOD levels in female flies of the positive group compared to the negative group suggest that the antioxidant response may be sex-specific and dose-dependent, as supported by Zhao *et al*. (2020), reported that female flies exhibit different oxidative stress responses compared to males. The higher SOD levels in females treated with 100 mg of MC indicate a possible protective effect at this dose. The significant increase in GSH levels in male flies of the positive group compared to the negative group is noteworthy. GSH is a critical antioxidant that plays a key role in detoxifying reactive oxygen species and maintaining redox balance. The decrease in MDA levels in female flies of the positive group suggests a reduction in lipid peroxidation, a common indicator of oxidative stress. This in line with research by Gohil *et al*. (2022), that antioxidant treatments can significantly reduce MDA levels, indicating a protective effect against oxidative damage. Conversely, the significant increase in MDA levels in male flies fed with 200 mg of MC indicates a potential pro-oxidant effect at higher doses, which has been documented in previous studies where high concentrations of phytochemicals led to increased oxidative stress (Mansoor *et al*., 2023). Female flies fed with 100 mg of MC exhibited the lowest MDA levels among the treatment groups, reinforcing the idea of a dose-dependent response.

The observed decrease in CAT levels in the positive control group males, particularly those treated with 200 mg of MC, suggests an impairment in the antioxidant defense mechanism, which aligns with findings by Zhang *et al*. (2022), observed that excessive oxidative stress can overcome antioxidant enzymes, reducing their activity. In contrast, the significant increase in CAT levels in the negative group of female flies indicates an adaptive response to maintain oxidative balance. This observation is critical, as maintaining CAT levels is essential for cellular protection against oxidative damage (Agarwal *et al*., 2021). The evident increase in SOD levels in the positive group male flies aligns with the findings of (Rojas *et al*., (2019), where oxidative stress led to the upregulation of SOD as a compensatory mechanism. However, the reduced SOD levels in female flies of the positive group compared to the negative group suggest that the antioxidant response may be sex-specific and dose-dependent, as supported by Zhao *et al*. (2020), reported that female flies exhibit different oxidative stress responses compared to males. The higher SOD levels in females treated with 100 mg of MC indicate a possible protective effect at this dose. The highest GSH levels observed in male flies fed with 100 mg of MC suggest that this dosage may enhance the antioxidant capacity effectively. Conversely, the low GSH levels in female flies treated with 150 mg and 200 mg of MC emphasize the potential for higher doses to disrupt the antioxidant defense mechanism, corroborating findings by Singh *et al*. (2021), reported that excessive oxidative stress can lead to GSH depletion in cellular systems. The highest GSH levels observed in male flies fed with 100 mg of MC suggest that this dosage may enhance the antioxidant capacity effectively.

The two strains of *D. melanogaster* (Harwich and indigenous) in this study exhibit different gene expression responses to both Metformin and MC, indicating genetic variability and nutrigenomic that influences glucose transporter regulation. While the Harwich strain shows a positive response to both Metformin and higher concentrations of MC, the indigenous strain shows a more significant negative response, particularly at higher concentrations of *M. charantia*. Tiannan, *et al*. (2020) mentioned that the differential response between strains highlights the importance of genetic background in determining the effectiveness of antidiabetic treatments, similar to findings in previous animal model studies. Metformin is a first-line therapy for the treatment of T2D due to its robust glucose-lowering effects, well-established safety profile. Relatively low cost and metformin has been shown to have pleiotropic effects on glucose metabolism, there is a general consensus that the major glucose-lowering effect in patients with type 2 diabetes is mostly mediated through inhibition of hepatic gluconeogenesis (Traci and Gerald 2021). Metformin appears to upregulate glucose transporter expression in both strains, though to a lesser extent in indigenous. Which is also stated that Metformin’s known mechanism involves the activation of AMP-activated protein kinase (AMPK), which enhances insulin sensitivity and increases the expression of glucose transporters like GLUT-4 in peripheral tissues (Traci and Gerald, 2021). In contrast, MC has a more variable effect depending on the concentration and the strain, and this could suggests that MC may have potential as a glucose transporter modulator but is highly dose-dependent and possibly strain-specific. The different concentrations of MC produce varying effects on glucose transporter gene expression. In the Harwich strain, higher concentrations of *M. charantia* (200 mg) seem to have a more beneficial effect (upregulation), while lower concentrations lead to downregulation. However, in the indigenous strain, higher concentrations of MC lead to significant downregulation, indicating potential toxicity or inhibitory effects at these levels. The differential expression of glucose transporters in response to MC could be due to its bioactive compounds interacting with insulin signaling pathways differently in each strain. The glucose transporter gene (GLUT) plays a central role in mediating glucose uptake into cells, particularly in tissues where insulin regulates glucose homeostasis (Aqeel *et al*., 2021). Changes in the expression of this gene could provide insight into how effectively glucose is being transported, which is in line with the study of Tian-Nan *et al*., (2022). And Albaik *et al*., (2024) mentioned that effective glucose is being transportation, is critical in managing hyperglycemia in diabetes.

Molecular docking was performed to evaluate the interaction of key phytochemicals with the *Drosophila* glucose transporter homolog *CG4607*, a parallel to human GLUT6. Among the docked compounds, squalene demonstrated the highest binding affinity (−7.5 kcal/mol), followed by bis(3-methylbutan-2-yl) phthalate and phytol. In vivo experiments on rats with T2DM showed that both metformin and squalene groups exhibited minor pancreatic rupture on histopathology. These findings, combined with in silico results, provide evidence of squalene’s antidiabetic and antioxidant properties. These interactions suggest that squalene may influence glucose uptake by modulating *CG4607* activity, potentially enhancing glucose transport and insulin sensitivity. These in silico findings offer a molecular rationale for the biochemical and physiological improvements observed in vivo and are in agreement with literature on the insulin-sensitizing effects of similar terpenoids (Ragasa *et al*., 2014).

Additionally, bis(3-methylbutan-2-yl) phthalate, found to have the second highest binding affinity in this study, has a potential to treat diabetes, due to its binding affinity with the modelled *CG4607* protein involved in glucose and lipid metabolism. This suggests that bis(3-methylbutan-2-yl) phthalate may also be effective in ameliorating type 2 diabetes. This is consistent with the work of Enenebeaku *et al*. (2021), who reported similar properties of bis(3-methylbutan-2-yl) phthalate in the context of antimalarial activity.

Multiple Sequence Alignment (MSA), a computational technique used to identify regions of similarity (Wu *et al*., 2025), and phylogenetic analysis, which represents the evolutionary relationships based on genetic similarities and differences (Mu *et al*., 2024), provided deeper insights into the genetic impact of treatments on *CG4607* gene. The clustering of MC-treated groups according to dosage confirms a concentration-dependent genomic response. In contrast, greater genetic variations were observed in diabetic control groups (Harwich-Positive (HP) and Ngd3-positive (NP)) corroborateing previous reports linking hyperglycemia to genomic instability in glucose metabolism pathways (Munteanu *et al*., 2025). It is well established that prolonged hyperglycemia induces oxidative stress, leading to nucleotide modifications and sequence variations in key metabolic genes (Bak *et al*., 2025). This reinforce the hypothesis that diabetes not only disrupts glucose uptake but also induces genetic alterations in *CG4607*, potentially contributing to disease progression.

The differential mutation profiles between Harwich and Ngd3 strains suggest a genetic background influence on *M. charantia* efficacy. Ngd3-treated groups (N100, N150, N200) exhibited more pronounced genetic variations than Harwich-treated groups (H100, H150, H200), indicating that the Ngd3 strain may have a heightened genomic response to phytochemical modulation. A similar strain-dependent variation in glucose metabolism gene expression was previously reported in *Drosophila melanogaster* model of diabetes (Kazek *et al*., 2024). Highlights the importance of genetic background in determining the therapeutic efficacy of plant-based interventions (Cai *et al*., 2025).

The phylogenetic tree revealed the formation of three main clusters, reflecting the influence of both strain background (Harwich and Ngd3) and treatment conditions. Samples from the Harwich strain treated with varying doses of *M. charantia* (H100, H150, H200) formed a tight cluster, indicating a high degree of genetic similarity under treatment. Similarly, the NgD3 samples treated with *M. charantia* (N100, N150, N200) grouped into a separate, distinct cluster. The greater divergence between Harwich and NgD3 strains is consistent with findings by Vesa *et al*. (2024), who reported that different *Drosophila* strains exhibit distinct metabolic adaptations to high-glucose diets. Furthermore, the clustering of MC-treated groups (H100, H150, H200, N100, N150, N200) into distinct dose-dependent branches aligns with research indicating that MC bioactive compounds (charantin, vicine, polypeptide-p) regulate glucose transporter genes (Khalid *et al*., 2023).

Interestingly, Metformin-treated samples (H16 for Harwich and N16 for Ngd3) clustered moderately close to their respective *M. charantia*-treated groups but showed slight divergence, particularly in the Ngd3 strain. This suggests that Metformin induces more pronounced genetic distinctiveness in Ngd3 compared to Harwich. These findings are consistent with Sakura *et al*. (2024), who demonstrated that Metformin modulates glucose transporter gene expression via AMPK activation in *Drosophila*. The observed phylogenetic divergence between H16 and N16 further supports a strain-dependent regulatory mechanism of for Metformin treatment, as also reported in previous transcriptomic studies by Alnuaimi *et al*. (2024). This supports the hypothesis that MC exerts its antidiabetic effects through mechanisms distinct from Metformin (Szymczak-Pajor *et al*., 2022). While Metformin primarily targets AMPK and insulin signaling pathways (LaMoia and Shulman, 2021), *M.charantia* likely regulates glucose transporter genes through direct antioxidant and anti-inflammatory mechanisms (Baenas and Wagner, 2022).

A third cluster comprising the untreated negative controls (HN, NN) and positive controls (HP, NP) showed partial genetic overlap, suggesting shared genetic responses in the absence of *M. charantia* or Metformin intervention. Clearer genetic separation was observed in the Ngd3 strain following treatment, particularly with Metformin, further emphasizing the role of genetic background and therapeutic intervention in shaping genetic variation. The observed genetic differences in *CG4607* between control groups supports the hypothesis that glucose transport mechanisms vary between *Drosophila* populations, possibly due to differences in insulin signalling efficiency (Bak *et al*., 2025). The higher bootstrap values (54%, 59%) associated with moderate and high MC doses suggest a stable genetic modifications likely mediated by phytochemical interactions with insulin-like peptides in *Drosophila* (Hussain *et al*., 2022). Additionally, the phylogenetic proximity of H200 and N200 indicates that treatment with *M. charantia* promotes genetic stability across different strains. This stability may reflect a protective effect against excessive genetic variation, potentially supporting more consistent long-term glucose regulation, in agreement with Wu *et al*. (2018), who showed that MC modulates glucose gene expression in a concentration-dependent manner.

### Conclusions

*Momordica charantia* ethanolic leaf extract is rich in bioactive compounds such as alkaloids, saponins, steroids, terpenoids, which may exhibit insulin-like activities and help regulate blood glucose. Molecular docking identified squalene (-7.5 kcal/mol) and bis(3-methylbutan-2-yl) phthalate (-7.4 kcal/mol) strong candidates for oral antidiabetic therapy based on their high binding affinities to the *CG4607* gene. Glucose levels were significantly reduced in both Harwich (0.75 mmol/L) and NgD3 (0.9 mmol/L) strains of *D. melanogaster* following co-feeding with with 200 mg and 100 mg of *M. charantia*. In the harwich strain triglyceride levels were highest (58.2 mg/dL) in the negative control, in the Ngd3 strain significantly decreased at 150 mg (5.55 mg/dL). Higher doses of *M. charantia* impaired climbing ability (p<0.001) but enhanced heat tolerance in the fruit flies. *Mormodica charantia* at 200 mg treatment induced strain-specific regulation of *CG4607* expression and genetic stabilization with upregulation in Harwich (0.5656) and a downregulation in indigeneous strains (-5.79586). Phylogenetic analysis revealed three distinct clusters.

### Recommendation

i. Other parts of the *M. charantia* fruits and roots should be further studied for the antidiabetes efficacy
ii. Studies could be focus on the specific pathways well as assessing the long-term impact of MC treatment on neuromuscular function and overall health in hyperglycemic models
iii. The other glucose transporter and insulin resistance genes should be further studied with the other strains of *D. melanogaster* including wild caught to have a wide range insight of the genetic variation
iv. The squalene and bis(3-methylbutan-2-yl) phthalate compounds should be synthesized and check the efficacy in further studies

## Funding

Tetfund

## Notes

### Competing Interest Statement

The authors have declared no competing interest.

